# Stimulus-selective lateral signaling between olfactory afferents enables parallel encoding of distinct CO_2_ dynamics

**DOI:** 10.1101/2020.12.03.410571

**Authors:** Dhruv Zocchi, Emily S. Ye, Elizabeth J. Hong

**Affiliations:** Division of Biology & Biological Engineering, California Institute of Technology, Pasadena, CA

**Keywords:** sensory processing, primary afferents, olfactory receptor neuron, *Drosophila*, carbon dioxide, lateral interactions, antennal lobe, temporal filtering, stimulus dynamics

## Abstract

An important problem in sensory processing is how lateral interactions that mediate the integration of information across sensory channels function with respect to stimulus tuning. We demonstrate a novel form of selective crosstalk between specific olfactory channels that occurs between primary olfactory receptor neurons (ORNs). Neurotransmitter release from ORNs can be driven by two distinct sources of excitation, feedforward activity derived from the odorant receptor and lateral input originating from specific subsets of other ORNs. Consequently, levels of presynaptic release can become dissociated from firing rate. Stimulus-selective lateral signaling results in the distributed representation of CO_2_, a behaviorally important environmental cue that elicits spiking in only a single ORN class, in multiple olfactory channels. Different CO_2_-responsive channels preferentially transmit distinct stimulus dynamics, thereby expanding the coding bandwidth for CO_2_. These results generalize to additional odors and olfactory channels, revealing a subnetwork of lateral interactions between ORNs that reshape the spatial and temporal structure of odor representations in a stimulus-specific manner.

**One Sentence Summary:** A novel subnetwork of stimulus-selective lateral interactions between primary olfactory sensory neurons enables new sensory computations.

## INTRODUCTION

Sensory circuits share a common functional architecture in which many parallel processing channels, each tuned to a specific feature of the external world, carry signals into the brain. In olfactory circuits, these processing channels correspond to anatomically discrete synaptic compartments called glomeruli. All olfactory receptor neurons (ORNs) expressing a given odorant receptor (OR) protein convergently project to a single glomerulus, and each second-order projection neuron (PN) receives direct input from a single glomerulus (Buck, 1996; Vosshall and Stocker, 2007). This orderly sorting of olfactory information, in which each processing channel is principally defined by the chemical selectivity of its cognate odorant receptor, is a hallmark of olfactory circuit organization. However, most stimuli, including most odors, activate multiple receptors, and the output of each glomerulus reflects the integration of direct input, governed by the activity of its cognate odorant receptor, and of indirect lateral inputs, derived from odorant receptors in parallel pathways. A core problem in sensory processing is to understand this integration, particularly the extent to which lateral interactions may or may not depend on the feature selectivity of the sensory channels they interconnect (Isaacson and Scanziani, 2011; Laurent, 1999). This question is important because stimulus-specific interactions could serve as the substrate for many types of useful sensory computations, such as feature extraction, signal decorrelation, or logic gating.

The antennal lobe of the fruit fly, *Drosophila melanogaster*, is numerically compact, comprising only ∼50 distinct glomeruli (Vosshall and Stocker, 2007), and is optically, electrophysiologically, and genetically accessible. In addition, its circuit architecture is strongly homologous to that of the vertebrate olfactory bulb and other early sensory circuits such as the inner retina (Wilson and Mainen, 2006). These properties make the antennal lobe a useful model for investigating the basic circuit mechanisms of sensory processing. In *Drosophila*, the responses of the uniglomerular projection neurons (PNs) arise from the integration of direct feedforward excitation from ORNs with indirect lateral inputs, mediated by networks of local neurons (Wilson, 2013). Lateral crosstalk in antennal lobe processing occurs as early as the primary afferents themselves, where global GABAergic inhibition of presynaptic ORN terminals regulates the gain of olfactory input signals (Olsen and Wilson, 2008; Root et al., 2008). Additionally, a small network of excitatory local neurons mediates weak coupling across PNs themselves (Olsen et al., 2007; Shang et al., 2007; Yaksi and Wilson, 2010). However, in both cases, these lateral networks act via broad, predominantly nonselective interactions between glomeruli to globally scale circuit function according to the overall ongoing level of activity in the network (Hong and Wilson, 2015; Olsen et al., 2007; Silbering and Galizia, 2007; Silbering et al., 2008).

Whereas most odors broadly activate multiple ORs, some specialized odors are recognized only by a dedicated OR subtype, and so are thought to activate only a single class of ORNs (Haverkamp et al., 2018; Hildebrand and Shepherd, 1997). Such specialist odors are usually ethologically significant to the animal, serving as salient signals for danger, food, or reproduction (Andersson et al., 2015; Depetris-Chauvin et al., 2015). One well-studied example of such a specialist odor is CO_2_, an important but complex sensory cue for animals across diverse phyla (Guerenstein and Hildebrand, 2008; Jones, 2013); it can assume positive or negative value, depending on context (van Breugel et al., 2018; Faucher et al., 2006). For example, CO_2_ is a major byproduct of microbial fermentation of organic substrates, a primary food source for *Drosophila* (van Breugel et al., 2018; Faucher et al., 2006). However, elevated CO_2_ has also been proposed to be a stress signal emitted by conspecifics (Suh et al., 2004), or a signal of potential predators or parasites. Behavioral aversion to CO_2_ is mediated by a single chemoreceptor complex Gr63a/Gr21a, acting in the ab1C class of ORNs (Jones et al., 2007; Kwon et al., 2007; Suh et al., 2004). Aversion to very high concentrations of CO_2_ also involves the acid-sensitive ionotropic receptor, Ir64a (Ai et al., 2010). However, recent work has also demonstrated that under certain highly active behavioral states, such as flight or rapid walking, flies are attracted to CO_2_ (van Breugel et al., 2018; Wasserman et al., 2013). This attraction is mediated by a chemosensory pathway distinct from that which controls aversion (van Breugel et al., 2018), motivating a search for additional classes of CO_2_-responsive ORNs.

In the course of this search, we uncovered a previously undescribed form of lateral information flow between ORNs through which signals elicited by many odors, including CO_2_, are transmitted between specific olfactory glomeruli. This lateral information flow is distinct from previously described forms of lateral connectivity in that is occurs selectively and directionally between specific glomeruli, forming subnetworks of glomeruli that are preferentially connected. A consequence of this new form of lateral information flow is that neurotransmitter release from ORNs can be significantly more broadly tuned than OR-mediated spiking responses at the corresponding cell body. We demonstrate that these lateral signals significantly restructure both the spatial and temporal dynamics of neurotransmitter release from olfactory afferents to postsynaptic targets in the brain.

## RESULTS

### ORNs in multiple glomeruli respond to CO_2_

Using in vivo two-photon calcium imaging, we volumetrically imaged from ORN axon terminals in the antennal lobe of flies expressing the genetically encoded calcium indicator GCaMP6f (Chen et al., 2013) in all ORNs, while delivering 3 s pulses of CO_2_ (Figure 1A, Figure S1). In addition to responses in the known class of CO_2_-responsive ORNs (ab1C) targeting the V glomerulus (Couto et al., 2005; Jones et al., 2007; Kwon et al., 2007; Suh et al., 2004), we also observed reliable, stimulus-locked calcium responses in at least three additional glomeruli at varying imaging depths. Based on their anatomical characteristics and odor responses (de Bruyne et al., 2001; Couto et al., 2005; Hallem and Carlson, 2006; Laissue et al., 1999), we identified these additional glomeruli as DL1, DM1, and VA2 (Figure 1A, and see Methods), which correspond to three ORN classes not previously reported as being responsive to CO_2_ (de Bruyne et al., 2001; Suh et al., 2004). CO_2_-evoked ORN responses in all four glomeruli were completely abolished by amputation of the third segment of the antenna (data not shown), which houses 90% of ORNs in fly, indicating that the responses are olfactory in origin.

**Figure 1:**
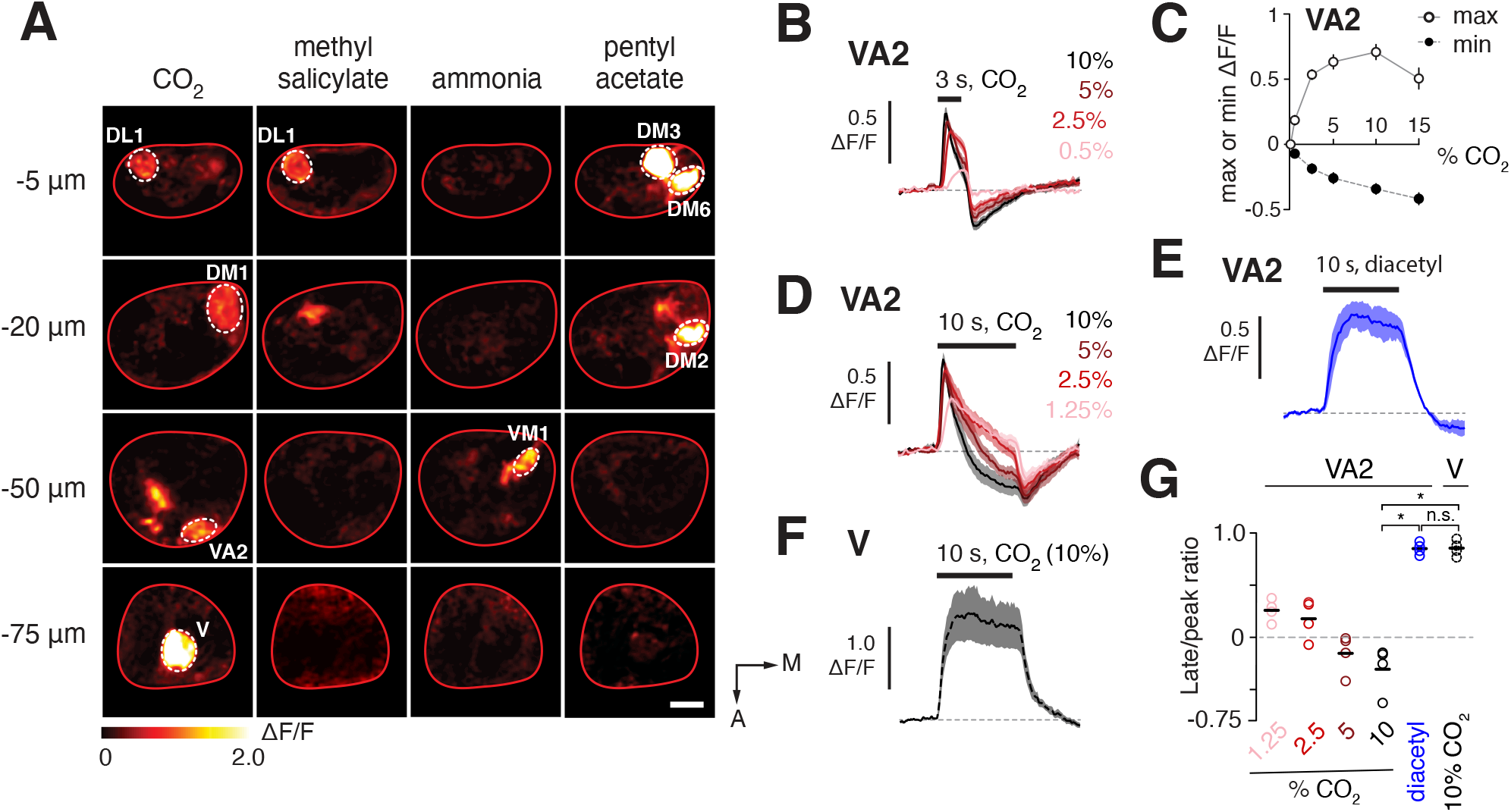
CO_2_ elicits mixed excitatory and inhibitory signals in multiple classes of ORNs. **A)** Two-photon imaging of odor-evoked calcium signals in ORN axon terminals in the antennal lobe. Images show representative peak *Δ*F/F responses from a fly expressing GCaMP6f under the control of the *pebbled-gal4* driver. The antennal lobe is viewed from above and is outlined in red (anterior is down, medial is right, scale bar is 20μm). Each row represents a different imaging plane, moving progressively deeper (more ventral) from the top to the bottom row. Dashed ovals indicate glomeruli activated by the odor panel and identified with high confidence based on location, size, shape, and odor tuning. CO_2_ responses were observed in ORNs in glomeruli DL1, DM1, VA2, and V. A subset of the odors most useful for identifying landmark glomeruli in each imaging plane are shown; they are 5% CO_2_, pentyl acetate (10^-5^), NH4 (10%), and methyl salicylate (10^-5^). **B)** Time course (mean and s.e.m.) of change in fluorescence in ORN terminals in VA2 to a 3 s pulse of CO_2_ at varying concentrations (*n*=3-9 flies, see Table S1). **C)** Maximum and minimum peaks (mean and s.e.m.) of the CO_2_ responses in **B**. **D)** Time course of *Δ*F/F response (mean and s.e.m.) of ORN terminals in VA2 to a 10 s pulse of varying concentrations of CO_2_ (*n* = 4 flies). **E)** Time course of *Δ*F/F response (mean and s.e.m.) of ORN terminals in VA2 to a 10 s pulse of diacetyl (10^-6^) (*n* = 4 flies). **F)** Time course of *Δ*F/F response (mean and s.e.m.) of ORN terminals in V to a 10 s pulse of 10% CO_2_ (*n* = 4 flies). **G)** Mean ratio (black bar) of the late response amplitude (defined as the response 8 s after the peak response) to the peak response amplitude in **D-F**. Each open circle is from an individual experiment. The late/peak ratio of CO_2_ responses becomes smaller as stimulus intensity increases (linear regression, R^2^ = 0.6053, p = 0.0004, slope = −0.0657 with 95% CI [-0.09612, −0.03529]). The late/peak ratio of the response to 10% CO_2_ of VA2 ORNs to significantly smaller than that of V ORNs, or of VA2 ORNs to diacetyl (\**p* < 0.05, repeated-measures ANOVA with Bonferroni multiple comparisons test; n.s. indicates not significantly different).

We initially focused on glomerulus VA2, which is innervated by the ab1B class of ORNs and easily identifiable due to its unique shape, position, and high sensitivity to diacetyl (de Bruyne et al., 2001; Couto et al., 2005; Fishilevich and Vosshall, 2005; Laissue et al., 1999) (Figure 1). VA2 axon terminals were sensitive to CO_2_ at concentrations ranging from 0.5% to 15% (Figure 1B-C) and exhibited unusual response dynamics, comprised of a fast excitatory component riding on a delayed inhibitory component, both of which scaled with stimulus intensity (Figure 1B-C). The excitatory component was more sensitive to CO_2_ but rapidly saturated as CO_2_ concentration increased, whereas CO_2_-evoked inhibition grew steadily with concentration (Figure 1B-C). The transient dynamics of the VA2 ORN response were even more apparent when we examined responses to a prolonged 10 s pulse of CO_2_ (Figure 1D). These responses were dominated by the inhibitory component, particularly for higher CO_2_ concentrations, and contrasted with the sustained and step-like response of VA2 ORN terminals to odors such as diacetyl or ethyl butyrate that elicit strong spiking responses in ab1B ORNs (Figure 1E, G; Figure 6B). Similarly, CO_2_-evoked responses in the V glomerulus, stemming from ab1C ORNs, also closely followed the shape of the stimulus, exhibiting a step-like, sustained profile (Figure 1F, G). The atypical temporal dynamics of CO_2_ signals in VA2 ORNs suggest they might arise through a different mechanism.

### Modulation of CO_2_-evoked responses in VA2

The results so far were surprising to us because prior studies using similar methods did not observe ORN responses to CO_2_ in any glomeruli besides V (Hong and Wilson, 2015; Suh et al., 2004). However, we found that the CO_2_-evoked responses in VA2 ORNs were unusual in that they were modulated over the duration of the recording. Early in each experiment, CO_2_ evoked a linearly increasing inhibition during stimulus presentation followed by a slightly faster recovery (Figure 2A-B, top row). At later time points, ORN activation by CO_2_ in VA2 transformed into a robust mixed response, comprised of rapid, phasic excitation at stimulus onset, followed by a delayed inhibition (Figure 2A-C). These experiments were conducted using 5% CO_2_, but similar results were observed with both lower and higher concentrations of CO_2_ (data not shown). These unusual results might be explained by changes in the health of the preparation during the experiment. However, concurrently measured CO_2_-evoked ORN responses in V did not change in amplitude throughout the full duration of the experiments. Similarly, ORN responses to diacetyl in VA2 were also stable, indicating that the potentiation of ORN responses to CO_2_ in VA2 is odor- and glomerulus-specific and thus cannot be easily explained by a general decline in the health of the preparation.

**Figure 2:**
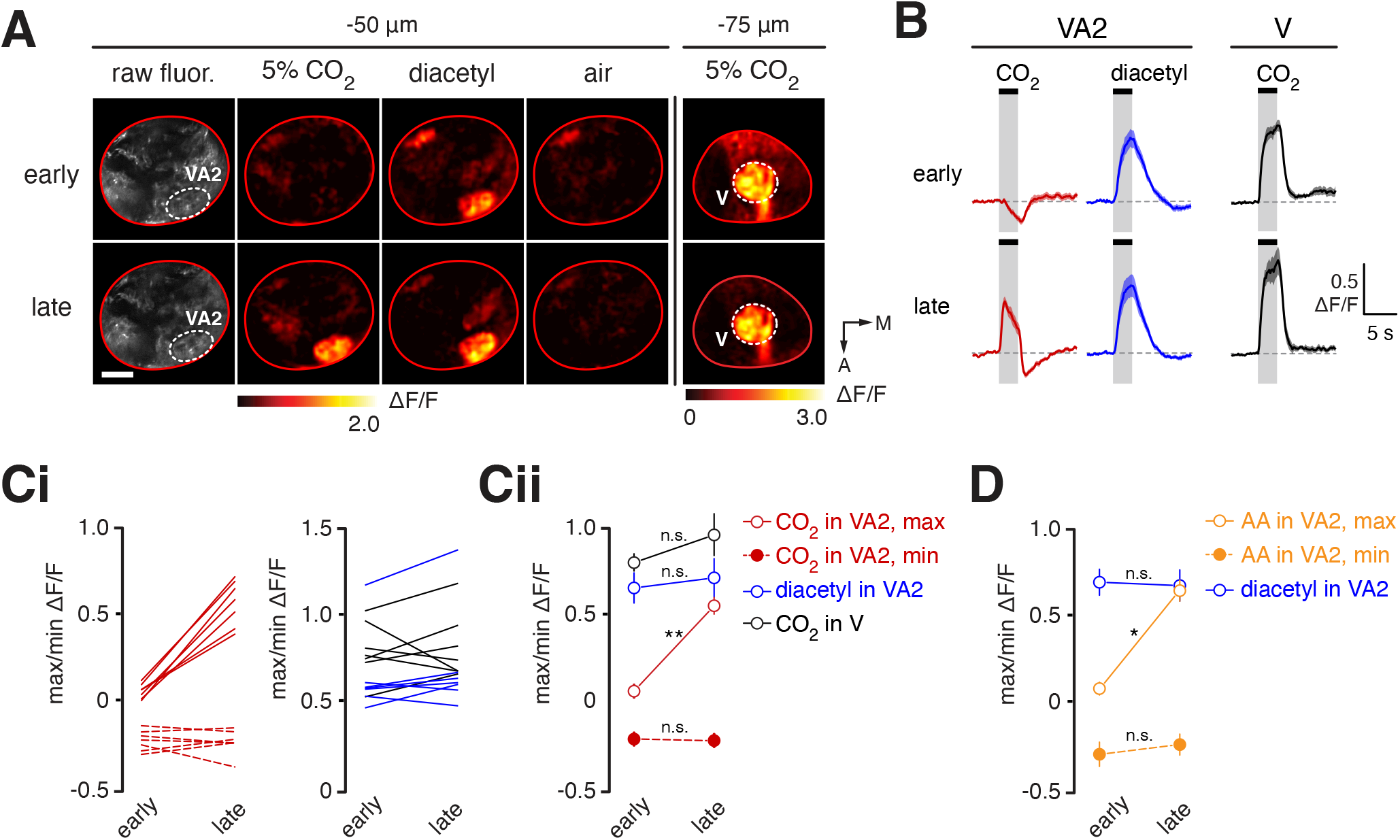
CO_2_ responses convert from a low to a high state over the course of a recording. **A)** Peak *Δ*F/F heat maps from a single experiment showing ORN responses to 5% CO_2_, diacetyl (10^- 6^), and control air in the two imaging planes containing glomeruli VA2 and V. The first column shows the resting fluorescence in the plane containing VA2, with the antennal lobe outlined in red (anterior is down, medial is right, scale bar is 20μm). The top row are responses recorded within the first 5 min of the experiment, and the bottom row shows responses in the same imaging field approximately 20 minutes later. **B)** Time course (mean and s.e.m.) of change in fluorescence in ORN terminals elicited by 5% CO_2_ (in VA2, red and V, black) and by diacetyl, 10^-6^ (in VA2, blue) at early and late time points during experiments (*n* = 7 flies). Early responses were from the first several presentations of each stimulus in an experiment, and late responses were defined as those collected >30 min after the start of the experiment and were stable in amplitude for multiple presentations of CO_2_ (see Method Details). **C)** Maximum (solid) and minimum (dashed) ORN responses in VA2 to 5% CO_2_ (red) or diacetyl, 10^-6^ (blue), and in V to 5% CO_2_ (black) at early and late time points. Maximum and minimum responses were determined within the 8 s window that starts at stimulus onset. Data from individual flies are shown in **Ci**, with early and late measurements from the same fly connected by a line. Mean maximum (open circles) and minimum (solid circles) responses (error bars are s.e.m.) are plotted in **Cii** (*n* = 7 flies). \*\**p*<0.01, two-tailed Mann-Whitney *U*-test with Bonferroni multiple comparisons test. **D)** Maximum (open circles) and minimum (solid circles) ORN responses (mean and s.e.m.) in VA2 to 3% acetic acid (orange) or diacetyl, 10^-6^ (blue) at early and late time points (*n* = 5 flies). Acetic acid-evoked responses of ORNs in VA2 also convert from a low to a high responsive state. \**p*<0.05, two-tailed Mann-Whitney *U*-test with Bonferroni multiple comparisons test; n.s. designates early/late comparisons that are not significant.

Quantification of VA2 ORN responses to CO_2_ across flies showed that the excitatory component of the response was selectively potentiated during the experiment whereas the inhibitory component remained stable (Figure 2C). This potentiation did not depend on prior exposure to CO_2_. Dissected experimental preparations that were left unstimulated for thirty minutes and then subjected to a single pulse of CO_2_ typically exhibited a full potentiated response in VA2 ORNs, comprised of mixed excitation and inhibition (data not shown). To examine the time course of the potentiation, we measured ORN responses in VA2 to a pulse of CO_2_ presented once every minute for 30 presentations (Figure S2A). The vast majority of flies (∼90%) started in the “low state” dominated by inhibition, and the excitatory component began to steadily increase in amplitude typically between ten and twenty minutes after the start of the experiment. Once the potentiation initiated, it typically continued unabated, and most flies exhibited the “high state” after thirty minutes. Although rare, we sometimes observed preparations that behaved differently, either starting the experiment already in the high state or remaining in the low state throughout (Figure S2B). Finally, we found that, although responses to most odors that activate VA2 were stable, responses of ORNs in VA2 to acetic acid, as well as to some additional short chain fatty acids, were also modulated over the course of the experiment, with characteristics mirroring those of CO_2_ (Figure 2D and data not shown). Thus, ORN responses in VA2 to CO_2_ and to some short chain fatty acids are modulated from a low to a high responsive state over the course of ∼30 min. This finding may account for why prior studies have not observed responses to CO_2_ outside the V glomerulus and further suggests that the response may arise via a separate mechanism from conventional ORN responses.

### CO_2_-evoked responses in VA2 occur in ab1B ORN terminals

We next investigated the origin of the CO_2_-evoked responses in VA2 ORN terminals. First, we revisited the question of whether VA2 ORNs fire action potentials in response to CO_2_. ORN cell bodies and their dendrites are grouped together in stereotyped combinations of 2-4 subclasses, with each grouping associated with a specific specialized sensory structure called a sensillum (Shanbhag et al., 1999, 2000). The ab1 sensillum houses four ORN subclasses, ab1A-D (Figure S3A). Prior studies surveying different ORN classes for CO_2_ sensitivity, as measured by single-sensillum extracellular recordings, did not observe spiking responses to CO_2_ in the ab1B class of ORNs (de Bruyne et al., 2001), which targets the VA2 glomerulus. In fact, only ab1C ORNs(de Bruyne et al., 2001), which target the V glomerulus, fired in response to CO_2_ (de Bruyne et al., 2001). We recorded extracellularly from ab1B ORNs and confirmed that they do not fire action potentials in response to CO_2_, at any concentration ranging from 0.5 to 10% CO_2_ (Figure 3A-B, and data not shown); ab1B ORNs also did not spike in response to acetic acid (Figure 3B). This experiment was performed in flies in which the ab1A ORN class was genetically ablated, to allow unambiguous identification of ab1B spikes (Figure 3A, Figure S3A-B). In order to also monitor calcium signals at ORN presynaptic terminals in this fly, we concurrently expressed GCaMP6f in many, but not all, ORN classes using the *orco* promoter (Couto et al., 2005) (Figure 3A). *Orco*, also known as *Or83b*, is a highly conserved co-receptor for insect ORs; it is expressed and required for olfactory transduction in ∼75% of the classes of ORNs, including ab1B (Larsson et al., 2004; Vosshall et al., 2000). Using these flies, we directly compared calcium responses in ORN terminals in VA2 using the same set of stimuli as for the single-sensillum recordings, and confirmed that, whereas neither CO_2_ nor acetic acid elicited spiking at ab1B ORN cell bodies, both odors evoked robust calcium responses in ORN axons in VA2 (Figure 3A-B). We note that, in single-sensillum recordings in which ab1B units was held for more than an hour, spiking responses to diacetyl and ethyl butyrate were stable throughout the duration of the experiment, whereas spiking responses to CO_2_ or acetic acid were never observed (data not shown).

**Figure 3:**
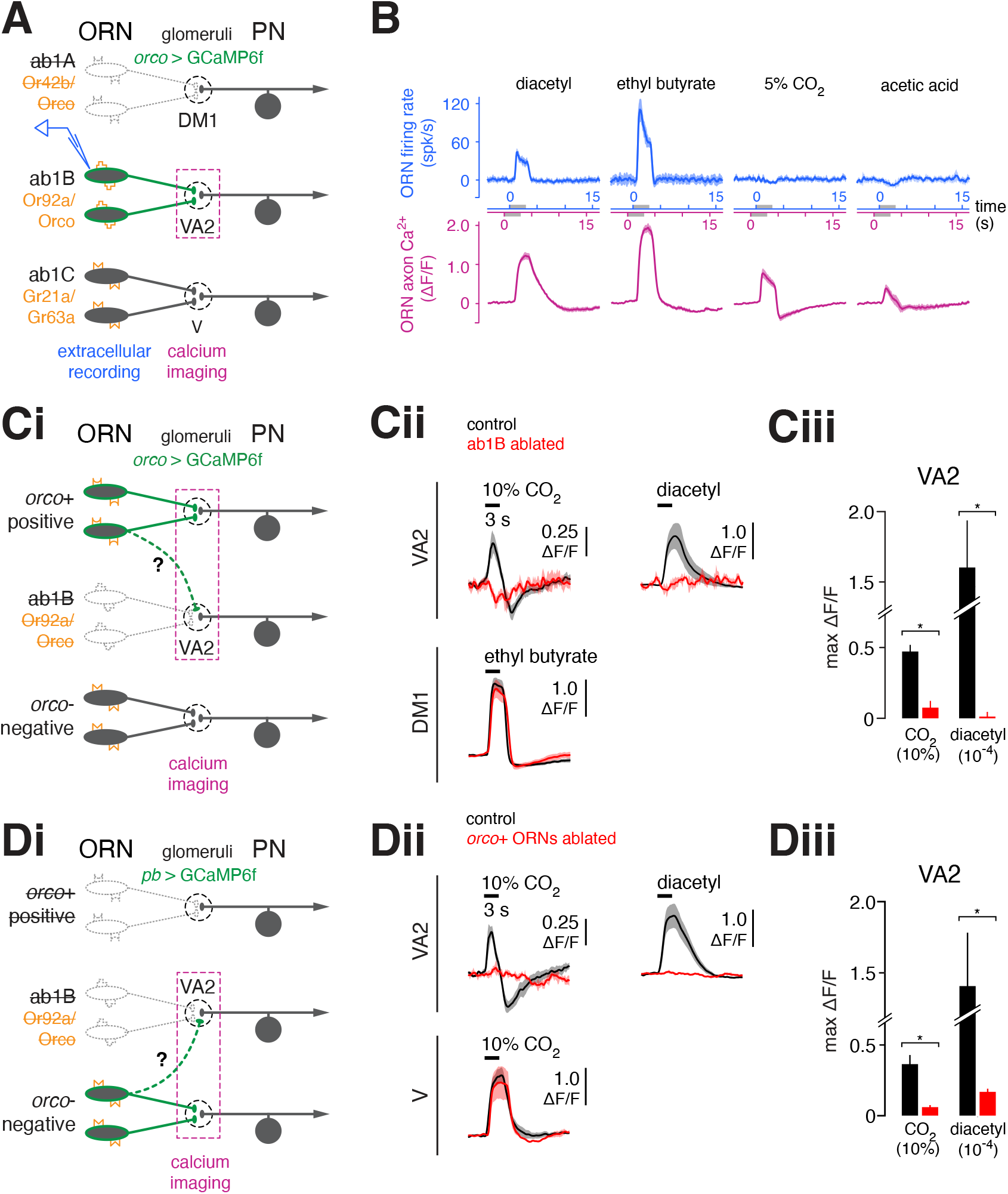
CO_2_-evoked calcium signals in VA2 occur in ab1B ORN terminals and are dissociable from ab1B spiking. **A)** Schematic of experimental setup. Odor-evoked action potentials in ab1B ORNs were measured by extracellular recordings targeting the ab1 sensillum. These single-sensillum recordings were conducted in flies in which the ab1A ORN (which has a similar spike amplitude to ab1B) was killed using diptheria toxin, to allow unambiguous identification of ab1B spikes (Figure S3A-B). Expression of GCaMP6f in from *orco-lexA* allowed imaging of odor-evoked calcium in ORN terminals in the same fly, as in Figure 1A. **B)** Comparison of odor-evoked responses in ab1B ORNs, as measured by spiking (blue) and presynaptic calcium in the axon terminal (magenta). The top row shows peristimulus time histograms (mean and s.e.m.) of odor-evoked ab1B firing (blue, *n* = 4-6 flies), and the bottom row shows the time course of *Δ*F/F response of ORNs in VA2 (magenta, *n* = 3-5 flies). Odors are diacetyl (10^-6^), ethyl butyrate (10^-4^), 5% CO_2_, and 3% acetic acid. **C-D)** CO_2_-evoked ORN calcium signals in VA2 occur in ab1B ORNs terminals. Stimuli are 10% CO_2_, diacetyl (10^-4^), and ethyl butyrate (10^-4^). **Ci)** Schematic of experimental setup. ab1B ORNs are ablated by expressing diphtheria toxin from *or92a*-*gal4*, while imaging ORN calcium in *orco*-positive ORNs. **Cii)** Odor-evoked calcium responses (mean and s.e.m.) from *orco*-positive ORNs in the indicated glomeruli in flies in which ab1B ORNs are ablated (red, *n* = 4 flies) or are intact (black, *n* = 7 flies). **Ciii)** Quantification of maximum peak odor-evoked *Δ*F/F responses (mean and s.e.m.) from **Cii**. \**p*<0.05, two-tailed Mann-Whitney *U*-test with Bonferroni multiple comparisons test. **Di)** Schematic of experimental setup. *orco*-positive ORNs are ablated by expressing diphtheria toxin from *orco*-*gal4*, while imaging ORN calcium from all remaining ORNs labeled by *pebbled*-gal4. **Dii)** Odor-evoked calcium responses (mean and s.e.m.) from ORNs in the indicated glomeruli in flies in which *orco*-positive ORNs are ablated (red, *n* = 3 flies) or are intact (black, *n* = 6 flies). **Diii)** Quantification of maximum peak odor-evoked *Δ*F/F responses (mean and s.e.m.) from **Dii**. \**p*<0.05, two-tailed Mann-Whitney *U*-test with Bonferroni multiple comparisons test.

We considered two possible explanations for the mismatch between the odor selectivity of ab1B ORN cell bodies and their presumptive axon terminals in VA2. The first possibility is that the CO_2_-evoked calcium response in VA2 does not, in fact, arise from ab1B axon terminals, but rather from a separate class of ORNs that also projects to VA2 (Figure 3Ci, Di). Although targeting of multiple glomeruli by a single ORN class is not known to occur in *Drosophila*, this model could account for our results, because the calcium indicator was driven from a genetic driver that labels every ORN in the circuit. The second possibility is that CO_2_-evoked calcium responses in VA2 arise from ab1B terminals but are driven by lateral input from another class of ORNs that directly detects CO_2_.

To distinguish between these possibilities, we investigated whether CO_2_-driven ORN calcium responses in VA2 occur in the ab1B terminals themselves. We expressed the A chain of diphtheria toxin (Han et al., 2000) (DTI) selectively in ab1B neurons using the promoter for its cognate odorant receptor, *or92a* (Couto et al., 2005), thereby killing this class of ORNs. Calcium signals in ORN terminals were monitored by placing GCaMP6f under the control of the *orco-gal4* transgene (Figure 3Ci). Ablation of ab1B neurons led to a complete loss of CO_2_-evoked responses in the VA2 glomerulus (Figure 3Cii-iii), demonstrating that responses to CO_2_ in VA2 are not arising from any non-ab1B, *orco*-positive ORNs with terminals in VA2. Other odors elicited robust calcium signals in *orco*-positive ORNs in other glomeruli (e.g. DM1, Figure Cii), demonstrating that the preparations were healthy and responsive. Next, to determine whether CO_2_-evoked responses in VA2 could arise from any *orco*-negative ORNs, we used DTI expression to ablate all *orco*-positive ORNs, which include the ab1B ORNs, and imaged calcium responses from all remaining *orco-*negative ORN terminals using a pan-ORN driver (Figure 3Di). Again, we observed no response to CO_2_ in VA2 (Figure 3Dii-iii), although responses to odor in glomeruli targeted by *orco-*negative ORNs (e.g. glomerulus V) were unaffected (Figure 3Dii). Because all ORNs are either *orco-*positive or *orco*-negative, these results collectively demonstrate that ORN calcium responses to CO_2_ in the VA2 glomerulus must occur in the terminals of ab1B ORNs.

### The excitatory and inhibitory components of CO_2_-evoked lateral input arise from genetically separable sources

These results suggest that CO_2_-evoked calcium responses in VA2 occur in ab1B terminals but are not driven by action potentials in ab1B ORNs. Thus, our data imply that ab1B axon terminals receive two distinct sources of excitation: direct excitation driven by Or92a-mediated olfactory transduction, and lateral excitation, which arises from one or more other ORN classes. To address this hypothesis, we tested whether CO_2_ responses in ab1B terminals in VA2 were indeed independent of OR function. Because a mutation in the *or92a* gene was not available, we instead functionally silenced ab1B ORNs, which are *orco*-dependent, by introducing a mutation in the *orco* gene (Larsson et al., 2004) in flies also expressing GCaMP6f in all ORNs (Figure 4A). We found that CO_2_-evoked responses in ab1B axon terminals in VA2 persisted in the *orco^-/-^* mutant (Figure 4Bi-ii). However, responses of ab1B terminals in VA2 to diacetyl, an odor that normally elicits robust firing in ab1B cell bodies (Figure 3B), were abolished in this fly, consistent with a loss of olfactory transduction in ab1B neurons (Fig 4Bi-ii). As expected, V glomerulus responses to CO_2_ in the axon terminals of ab1C ORNs, which are not *orco*-dependent, were unaffected (Fig 4Bi-ii). We noted, however, that the dynamics of the CO_2_ response in VA2 were fundamentally altered in the *orco^-/-^* mutant as compared to wildtype flies (Figure 4Bi); whereas the excitatory component remained, the inhibitory component was eliminated. This observation indicates that CO_2_-evoked inhibition in ab1B terminals depends on the activity of an *orco*-dependent OR(s). Collectively, these results demonstrate that CO_2_-evoked excitation in ab1B axon terminals is not driven by olfactory transduction in ab1B ORNs. Furthermore, they imply that ab1B ORNs receive an *orco-*independent source of lateral excitatory input at some location between the cell body and the presynaptic terminal.

**Figure 4:**
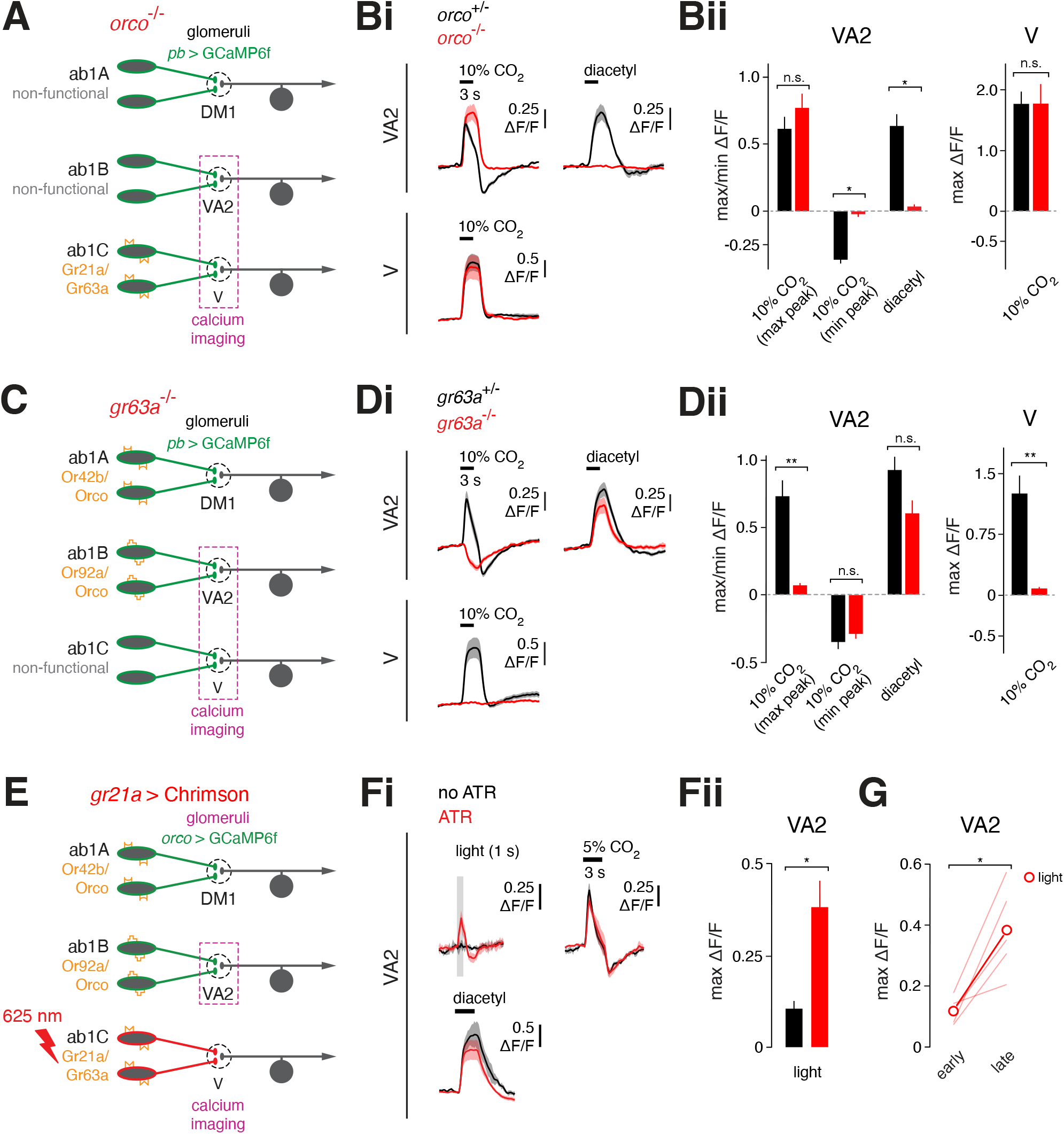
CO_2_-evoked calcium signals in VA2 are driven by lateral input from distinct, genetically separable sources. **A)** Schematic of experimental setup. Odor-evoked calcium responses are measured in *orco*^-/-^ flies expressing GCaMP6f in all ORNs (from *pebbled-gal4*). The loss of Orco abolishes odor transduction in all *orco-*positive ORNs, which include ab1B. Odor stimuli are 10% CO_2_ and diacetyl (10^-6^) in all experiments in this figure, except for panels **Fi-ii** in which 5% CO_2_ is used. **Bi)** Odor-evoked calcium responses (mean and s.e.m.) of ORNs in glomeruli VA2 and V in heterozygous control (black, *n* = 4 flies) and *orco*^-/-^ null flies (red, *n* = 7). **Bii)** Quantification of maximum/minimum peak odor-evoked *Δ*F/F responses (mean and s.e.m.) from **Bi**. \**p*<0.05, two-tailed Mann-Whitney *U*-test with Bonferroni multiple comparisons test. n.s., not significant. **C)** Schematic of experimental setup. Odor-evoked calcium responses were measured in *gr63a*^-/-^ null flies expressing GCaMP6f in all ORNs (from *pebbled-gal4*). The loss of *gr63a* abolishes odor transduction in ab1C ORNs. **Di)** Odor-evoked calcium responses (mean and s.e.m.) of ORNs in glomeruli VA2 and V in heterozygous control (black, *n* = 6 flies) and *gr63a*^-/-^ null flies (red, *n* = 7). **Dii)** Quantification of maximum/minimum peak odor-evoked *Δ*F/F responses (mean and s.e.m.) from **Di**. \*\**p*<0.01, two-tailed Mann-Whitney *U*-test with Bonferroni multiple comparisons test. n.s., not significant (*p*>0.05). **E)** Schematic of experimental setup. Calcium responses are measured from ORNs in VA2 while optogenetically exciting ab1C ORNs. Flies express GCaMP6f under the control of *orco-lexA* and CsChrimson under the control of *gr21a-gal4*. **Fi)** Calcium responses (mean and s.e.m.) of ORNs in VA2 evoked by delivering a 1 s pulse of light (625 nm) to flies raised on food without (control, black, *n* = 3 flies) or supplemented with (ATR, red, *n* = 4-5 flies) all-*trans*-retinal. Odor-evoked responses of ORNs in VA2 to CO_2_ and diacetyl are also shown for comparison. **Fii**) Quantification of the maximum peak of light-evoked *Δ*F/F responses (mean and s.e.m.) from **Di**. \**p*<0.05, two-tailed Mann-Whitney *U*-test. **G)** Responses of ORNs in VA2 evoked by light-based activation of ab1C convert from a low to a high response state. Maximum peak light-evoked ORN responses in VA2 from individual flies (*n* = 5 flies) raised on ATR at early and late times in the experiment, as defined in Figure 2B. Measurements from each individual fly are connected with a light red line. The bold open symbols are the mean response at early and late time points. \**p*<0.05, two-tailed Mann-Whitney *U*-test.

We next sought to identify the source for CO_2_-driven lateral excitation in ab1B ORNs. The principal sensor for CO_2_ in flies is the Gr63a/Gr21a receptor complex in ab1C ORNs, which project to glomerulus V (Jones et al., 2007; Kwon et al., 2007; Scott et al., 2001). To test whether the Gr63a/Gr21a receptor complex is required for CO_2_-driven lateral excitation to ab1B neurons, we introduced a mutation in *gr63a* (Jones et al., 2007) into flies also expressing GCaMP6f in all ORN classes (Figure 4Ci). This mutation renders the Gr63a/Gr21a complex non-functional (Jones et al., 2007), as confirmed by the complete loss of CO_2_-evoked firing in ab1C ORNs (data not shown). When we examined odor responses in ab1B axons in VA2 using calcium imaging, we found that CO_2_-evoked excitation was absent in *gr63a*^-/-^ flies, whereas axonal responses to diacetyl, which directly elicits spiking in ab1B ORNs (Figure 3B), were intact (Figure 4Di-ii). CO_2_-evoked calcium signals in the V glomerulus were completely absent, confirming that ab1C ORNs were silenced. Further, although the excitatory component was absent from ab1B responses to CO_2_ in VA2, the inhibitory component persisted (Figure 4Di-ii). Taken together, these results demonstrate that the excitatory and inhibitory components of CO_2_-driven lateral input arise from distinct receptors, and the source of CO_2_-driven lateral excitation in ab1B ORNs is the Gr63a/Gr21a receptor complex.

We reasoned that if CO_2_-driven excitation in ab1B terminals in VA2 depends on Gr63a/Gr21a function in ab1C neurons, then direct optogenetic stimulation of ab1C ORNs should be sufficient to evoke calcium signals in ab1B terminals. To test this, we used the *gr21a-gal4* driver (Couto et al., 2005) to express the red-shifted channelrhodopsin CsChrimson (Klapoetke et al., 2014) in ab1C ORNs, while simultaneously expressing GCaMP6f in a broad set of ORNs under the control of the *orco* promoter (Figure 4E). We then imaged ORN terminals in VA2 while stimulating the antennae with red light (625 nm). Light-driven activation of ab1C ORNs (Figure S3C) evoked calcium responses in ab1B terminals in VA2, and these responses were dependent on raising the flies on retinal, which serves as the chromophore for channelrhodopsin (Figure 4Fi-ii). Like the responses elicited by CO_2_ stimulation, ab1B responses driven by optogenetic activation of ab1C ORNs also changed over time, increasing in amplitude during the course of the experiment (Figure 4G). These results directly demonstrate that excitation in ab1B terminals originates from neural activity in ab1C ORNs and does not stem from direct or indirect effects of CO_2_ on a secondary target.

We next investigated the synaptic mechanisms by which ab1B ORNs receive lateral input. The major excitatory neurotransmitter in the antennal lobe is acetylcholine (Kazama and Wilson, 2008; Wilson et al., 2004). CO_2_-evoked calcium signals in ORNs in VA2 were undiminished by application of the nicotinic acetylcholine receptor antagonist mecamylamine (Kazama and Wilson, 2008; Wilson et al., 2004) (Figure S4Ai-ii). However, in separate experiments in which we imaged responses in PNs, mecamylamine was only partially effective in blocking calcium responses in PN dendrites arising from direct ORN input (Figure S4Ai-ii). Together, these results suggest that CO_2_-evoked lateral input to ab1B ORNs does not strongly depend on cholinergic transmission, but they cannot rule out a partial contribution. We next applied a cocktail of GABA receptor antagonists (Wilson and Laurent, 2005) to flies expressing GCaMP6f in all ORNs and observed a large increase in the baseline fluorescence in ab1B ORN terminals (Figure S4Bii), indicating they normally experience strong tonic inhibition. With GABA receptor blockade, odor-evoked responses in ab1B ORN terminals were present but significantly blunted, likely because of saturation of the calcium indicator. This effect was observed for both lateral signals evoked by CO_2_, as well as for conventional calcium responses evoked by odors that elicit direct excitation (Figure S4Bi). When we increased the concentration of the GABAA-receptor antagonist picrotoxin to levels at which inhibitory glutamatergic signaling is also impaired (Liu and Wilson, 2013), we observed similar results (data not shown). These results indicate that neither GABAergic nor glutamatergic function is required for CO_2_-evoked lateral signaling to ab1B ORNs.

Having not identified a strong contribution from the major classes of chemical neurotransmitters in the antennal lobe, we considered additional modes of intercellular signaling. For instance, prior work shows that the gap junction subunit shakB is required for electrical coupling in the optic lobe, in the giant fiber pathway, and for lateral excitatory interactions between PNs in the antennal lobe (Phelan, 2005; Yaksi and Wilson, 2010). We introduced the *shakB^2^* mutation (Krishnan et al., 1993) into flies expressing GCaMP6f in all ORNs and found lateral inputs to ab1B ORNs were unaffected in *shakB^2^* flies (Figure S4C). Finally, the Na^+^/K^+^-ATPase pump has been previously implicated in non-canonical cell-cell interactions (Kehoe and Ascher, 1970; Pinsker and Kandel, 1969; Pivovarov et al., 2018). Application of the Na^+^/K^+^-ATPase inhibitor ouabain selectively abolished the excitatory component of CO_2_-evoked responses in ab1B terminals (Figure S4D). Odor responses stemming from direct ORN activity were unaffected. These results were unexpected, since any direct electrogenic action of the Na^+^/K^+^-pump would be expected to contribute to the inhibitory component of the response. However, suppression of Na^+^/K^+^-pump function has many secondary consequences, for instance, changes in intracellular Ca^2+^ that can modulate membrane receptor and gap junction function (Matchkov Vladimir V. et al., 2007; Pivovarov et al., 2018; Weingart, 1977). Future work will be required to determine the mechanism by which ouabain blocks CO_2_-evoked lateral excitation in ab1B axon terminals.

### Lateral information flow among additional classes of ORNs

The lateral information flow to ab1b neurons might represent a phenomenon specific to that ORN class, or alternatively, might occur in other ORNs. To address this possibility, we examined a second class of ORNs – the ab1A ORNs which project to glomerulus DM1. We compared odor tuning at the cell body, as measured by extracellular recordings, to that at the axon terminal, as measured by calcium imaging (Figure 5A). As before, we observed differences in odor-evoked spiking and presynaptic calcium in DM1 ORNs (Figure 5B, Figure S5A-B). Most notably, CO_2_ and pentanol, odors that did not elicit changes in ab1A ORN firing rates, elicited robust calcium signals in the axon terminals. Due to the convergence of dozens of axons from ORNs with the same receptive field in a single glomerulus (de Bruyne et al., 2001; Vosshall et al., 2000), calcium imaging of population axonal responses in a glomerulus could feasibly achieve greater sensitivity than extracellular recordings from single neurons averaged across only a few trials. To directly evaluate whether differences in measurement methodologies, rather than lateral ORN input, underlie the difference between odor-evoked spiking and presynaptic calcium signals, we abolished olfactory transduction in ab1A ORNs (Figure 5C). This experiment was possible in ab1A ORNs, but not ab1B ORNs, because a null mutant in the ab1A odorant receptor *or42b* was readily available, which was not the case for *or92b*, the odorant receptor in ab1B ORNs. Whereas odor-evoked firing at ab1A ORN cell bodies was completely eliminated in *or42b*^-/-^ flies (Bellen et al., 2011) (Figure 5D, blue traces), ab1A terminals still responded robustly to odor (Figure 5D, magenta traces, Figure S5C-D). For most odors, the presynaptic Δ*F*/*F* response was comparable or larger when the OR was mutated, and the rank order preference for odors was also changed (Figure 5D). A notable exception was ethyl acetate which, at the concentrations used in this study, is known to selectively activate ab1A ORNs only (Olsen et al., 2010) (Figure S5B). Ethyl acetate (10^-8^) did not evoke calcium responses in ab1A terminals of *or42b*^-/-^ flies (Figure 5D, Figure S5D), consistent with its inability to activate any non-ab1A ORNs that could serve as sources of lateral input. Like CO_2_ responses in ab1B ORNs, laterally derived CO_2_ responses in ab1A ORNs were comprised of an excitatory and an inhibitory component, although the inhibitory component was comparatively less prominent (Figure 5D). These results show that, like ab1B ORNs, ab1A ORNs receive lateral input from ab1C ORNs, as well as many other classes of ORNs activated by the odors in our stimulus panel.

**Figure 5:**
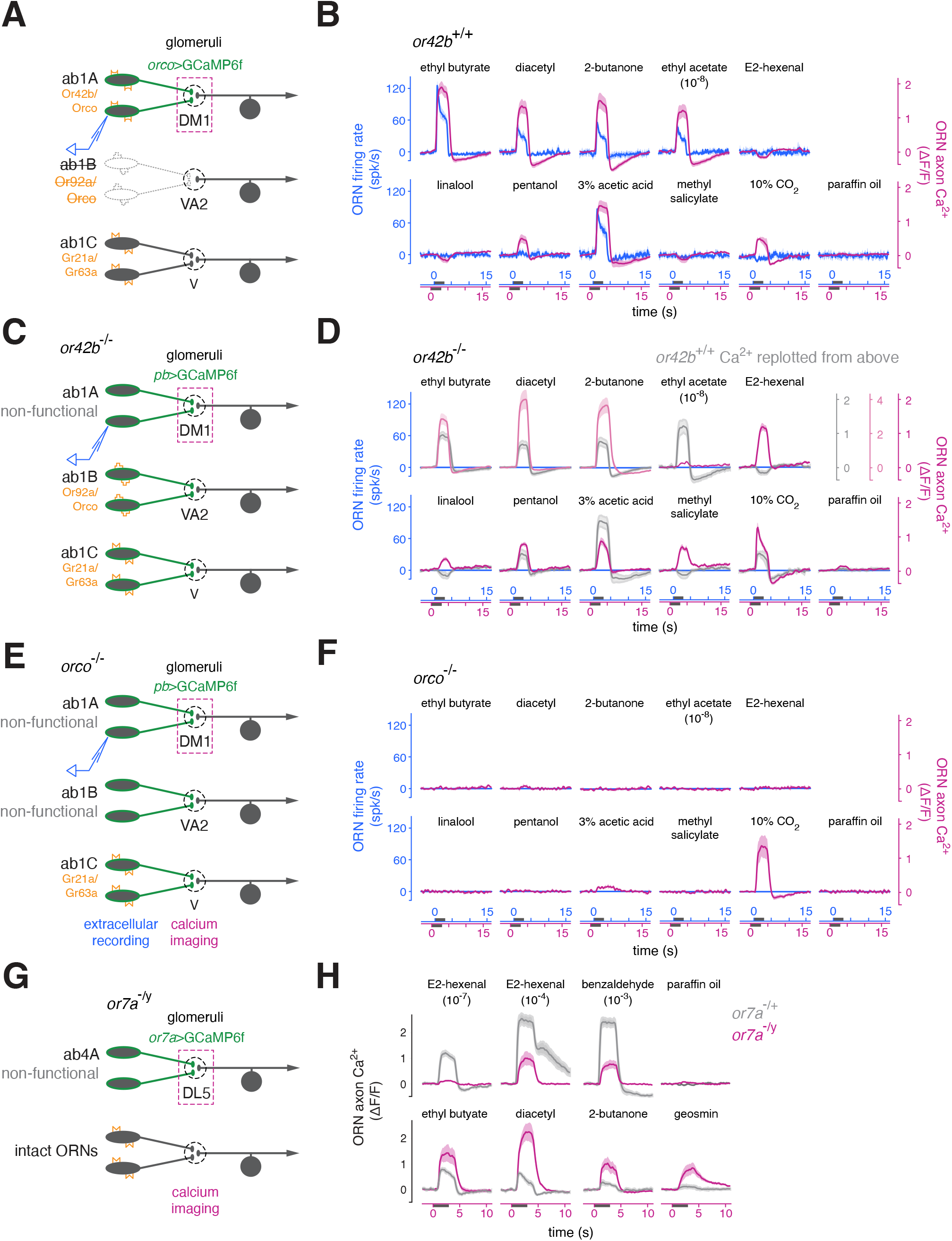
Odor-evoked activity in the axon terminals of functionally silent ab1A ORNs demonstrates selective lateral input to ab1A ORNs from other ORN classes. **A)** Schematic of experimental setup. Odor-evoked action potentials in ab1A ORNs were measured by ab1 sensillum recordings. These recordings were performed in flies in which the ab1B ORN was ablated using diptheria toxin, to allow unambiguous identification of ab1A spikes (Figure S3A-B). Expression of GCaMP6f from *orco-lexA* allowed imaging of calcium signals in ORN terminals in the same fly. **B)** Comparison of peristimulus time histograms (mean and s.e.m.) of odor-evoked ab1A spiking responses (blue, *n* = 3 flies) and the time course of odor-evoked change in fluorescence in ab1A axon terminals (magenta, *n* = 4-11 flies). For panels B, D, and F, the time axes for calcium responses are shifted forward slightly relative to that of the firing rate responses for visualization; the shift aligns the onset of the two types of responses due to the slower kinetics of the calcium signal. All odors were presented at a dilution of 10^-4^, with the exception of ethyl acetate (10^-8^) and CO_2_ (10%) **C)** Schematic of experimental setup. The *or42b^-/-^* mutation abolishes odor transduction in ab1A ORNs. Expressing GCaMP6f in all ORNs (from *pebbled-gal4*), and imaging from ab1A terminals in DM1, allows measurement of calcium responses in the axon terminals of ORNs which have no spiking responses to odor. **D)** Extracellular sensillum recordings (blue traces, *n* = 3 flies) confirm that ab1A ORNs in *or42b*^-/-^ mutant flies do not fire in response to any odor, but robust odor-evoked calcium responses persist in ab1A ORN terminals in DM1 in response to most stimuli (magenta traces, *n* = 6-13 flies). Odor-evoked calcium responses from ORNs in DM1 in wildtype flies are overlaid in grey for comparison. **E)** Schematic of experimental setup. Odor-evoked calcium responses are measured in *orco*^-/-^ null flies expressing GCaMP6f in all ORNs (from *pebbled-gal4*). The *orco*^-/-^ mutation abolishes odor transduction in ∼75% of all ORN types, including ab1A. **F)** Extracellular sensillum recordings confirm that the *orco*^-/-^ mutation eliminates odor-evoked spiking in ab1A ORNs (blue traces, *n* = 3 flies). Loss of odor transduction in additional classes of *orco*-dependent ORNs leads to a loss of nearly all laterally derived odor-evoked calcium responses (mean and s.e.m., *n* = 3-5 flies) in ab1A terminals in DM1 (compare to D). The exception is the lateral response evoked by CO_2_, which directly excites the *orco*-independent ab1C ORN class. **G)** Schematic of experimental setup. Knock-in of *gal4* into the endogenous *or7a* locus simultaneously disrupts the function of Or7a and facilitates expression of GCaMP6f selectively in ab4A ORN terminals targeting the DL5 glomerulus. H) Odor-evoked calcium responses in ab4A ORN terminals in *or7a*^-/y^ hemizygous (magenta, *n* = 4-5 flies) and *or7a*^-/+^ heterozygous (control, grey, *n* = 4-6 flies) flies. ab4A ORNs in *or7a*^-/y^ hemizygous flies lack a functional odorant receptor. Calcium responses to odors that directly activate ab4A ORNs (E2-hexenal, benzaldehyde) were diminished in the absence of *or7a* function compared to controls, as expected. All PSTH and calcium response time courses are plotted as mean and s.e.m. in all panels. Odors were presented at a dilution of 10^-4^, except where otherwise indicated. Like in ab1B ORNs, lateral signals in ab1A and ab4A ORN terminals modulate over time; all recordings in the figure were performed >30 min from the start of the experiment.

One important caveat is that odor-evoked lateral responses in DM1 in *or42b*^-/-^ mutants might reflect the misidentification of the DM1 glomerulus or the detection of strong out-of-plane signals due to imperfect optical sectioning during two-photon excitation. To confirm that the calcium signals we measured were from ab1A ORNs, we imaged from DM1 in flies in which we ablated ab1A ORNs using diptheria toxin, while expressing GCaMP6f from the *orco* promoter (which labels ab1A ORNs if present) (Figure S5E). The baseline fluorescence in DM1 was noticeably absent from these flies, and odor-evoked signals were absent at the expected position of the DM1 glomerulus (Figure S5F). Next, we imaged again from DM1 in *or42b*^-/-^ mutant flies, but we restricted the expression of GCaMP6f to ab1A ORNs, using the *or42b-gal4* driver (Fishilevich and Vosshall, 2005) (Figure S5G). As before, we observed robust odor-evoked calcium signals in ab1A presynaptic terminals in response to many odors (Figure S5H). In this experiment, we were unambiguously recording calcium responses from the axon terminals of ab1A ORNs completely lacking olfactory transduction and odor-evoked spiking responses; therefore, we conclude that these ORNs must be receiving lateral inputs from other classes of ORNs.

If calcium responses in ab1A presynaptic terminals arise from activity in other ORNs, silencing other ORN classes in addition to ab1A should reduce these responses. Odor-evoked calcium signals in DM1 ORN terminals were nearly eliminated in the *orco*^-/-^ mutant, with the exception of responses elicited by CO_2_ (Figure 5F, magenta traces). This observation is consistent with the fact that CO_2_ is detected by *orco-*independent Gr21a/Gr63a-expressing ORNs. We noted that, like in VA2, the responses of DM1 ORN terminals to some odors, including CO_2_, had a pronounced inhibitory component (Figure 5B). The inhibitory component of the CO_2_-evoked response is not due to direct suppression of odorant receptor activity, since it persisted in the *or42b*^-/-^ mutant (Figure 5D) but was substantially reduced in *orco*^-/-^ flies. Thus, like ab1B ORNs, ab1A ORNs also receive lateral inhibitory input stemming from activity in *orco*-positive ORNs; this inhibition is distinct from the well-described form of GABAergic presynaptic inhibition at ORN terminals, because it is insensitive to GABA receptor antagonists (Figure S4B and data not shown).

Finally, we investigated a third class of ORNs, the ab4A ORNs expressing the Or7a odorant receptor and projecting to the DL5 glomerulus (Couto et al., 2005; Vosshall and Stocker, 2007). ab4A ORNs are housed in a different olfactory sensillum on the antenna from ab1A and ab1B ORNs. We used a fly in which the *gal4* gene was knocked-in to the *or7a* locus, disrupting *or7a* function and selectively expressing Gal4 in ab4a ORNs (Figure 5G, (Lin et al., 2015)). We examined responses in ab4A ORN terminals to both odors that strongly activate ab4A ORN spiking (E2-hexenal and benzaldehyde) and to odors that do not elicit spiking activity in ab4A ORNs (ethyl butyrate, 2-butanone, and geosmin) ((Hallem and Carlson, 2006), and data not shown). Stimulation of *or7a*^-/y^ flies with this diverse panel of odors elicited large calcium responses in the terminals of ab4A ORNs, which lacked their normal odorant receptor Or7a (Figure 5H). As expected, stimuli that directly excite the Or7a receptor strongly, such as E2-hexenal and benzaldehyde (Hallem and Carlson, 2006), elicited weaker calcium signals in terminals of ab4A ORNs without a functional odorant receptor compared to heterozygous *or7a*^-/+^ flies. For instance, calcium responses were mostly abolished in response to E2-hexenal (10^-7^), an odor stimulus that selectively elicits firing in ab4A ORNs and does not directly excite additional ORN classes that could be sources of lateral input to ab4A terminals (Hong and Wilson, 2015; Olsen et al., 2010). However, stimuli that broadly excite many ORs beyond Or7a elicited strong calcium responses in the terminals of ab4A ORNs in *or7a*^-/y^ flies; these responses must arise from odorant receptor activity in non-ab4A ORNs.

Taken together, these results demonstrate that ab1A and ab4A ORNs also receive lateral signals that restructure presynaptic calcium responses. Thus, stimulus-selective lateral interactions between ORNs are not just unique to ab1B cells, but rather reflect a more general feature of the network and extend beyond the processing of CO_2_ signals.

### Dual encoding of CO_2_ in different glomeruli with distinct response dynamics

Finally, we considered how this newly identified form of lateral input to ORN presynaptic terminals impacts the transmission of stimuli across ORN-PN synapses. In particular, we focused on the implications for the encoding of CO_2_ signals, which exhibit dynamics that vary across a wide range of timescales. For instance, while navigating natural odor plumes in flight, *Drosophila* may experience rapid fluctuations in CO_2_ emanating from a distant fermenting source (Murlis et al., 1992; van Breugel and Dickinson, 2014), whereas flies walking across the surface of a rotting fruit would encounter slow, local CO_2_ fluctuations. As shown earlier, when flies were presented with a sustained pulse of CO_2_, ORN terminals in VA2 (i.e., ab1B ORNs, and hereafter referred to as VA2 ORNs) exhibited a transient excitation that rapidly adapted below baseline; this adaptation was driven by a delayed inhibitory component of the lateral input that quickly dominated the response (Figures 1D, 6A). These strongly adapting, transient dynamics are unique to ORN presynaptic calcium arising from lateral input and contrast with the non-adapting presynaptic calcium signal stemming from conventional direct OR-mediated excitation in either VA2 ORNs (e.g. by diacetyl or ethyl butyrate, Figure 6B) or in V ORNs (i.e. ab1C ORNs driven by CO_2_) (Figures 1E-F). When we measured calcium signals in the dendrites of VA2 PNs, which receive direct input from VA2 ORNs, we found that the strongly adapting dynamics of VA2 ORN calcium responses to CO_2_ were largely inherited by postsynaptic PNs. VA2 PNs responded phasically at the onset and the offset of the CO_2_ pulse, encoding rapid changes in the concentration of CO_2_ (Figure 6C). In contrast, VA2 PNs responded tonically to diacetyl or ethyl butyrate (Figure 6D and data not shown), more faithfully following the absolute levels of the stimulus. Similar results were also observed in DM1 ORNs and PNs, although DM1 PNs responded to the offset of the CO_2_ stimulus with a transient hyperpolarization rather than depolarization (Figure S6). When we examined responses in VA2 and DM1 to 1 Hz trains of CO_2_ or ethyl butyrate, ORNs were able to follow the individual pulses of either odor, but encoded CO_2_ fluctuations with comparatively higher contrast (Figure 6A-B, Figure S6A-B). PN calcium showed a similar, though less pronounced, effect (Figure 6C-D, Figure S6C-D). Together these findings show that the same ORNs can impose very different temporal filters on odor stimuli depending on their chemical identity, that these dynamics shape downstream responses in PNs, and that one mechanism by which this can occur is through the interplay of lateral excitatory and inhibitory inputs at ORN terminals.

**Figure 6:**
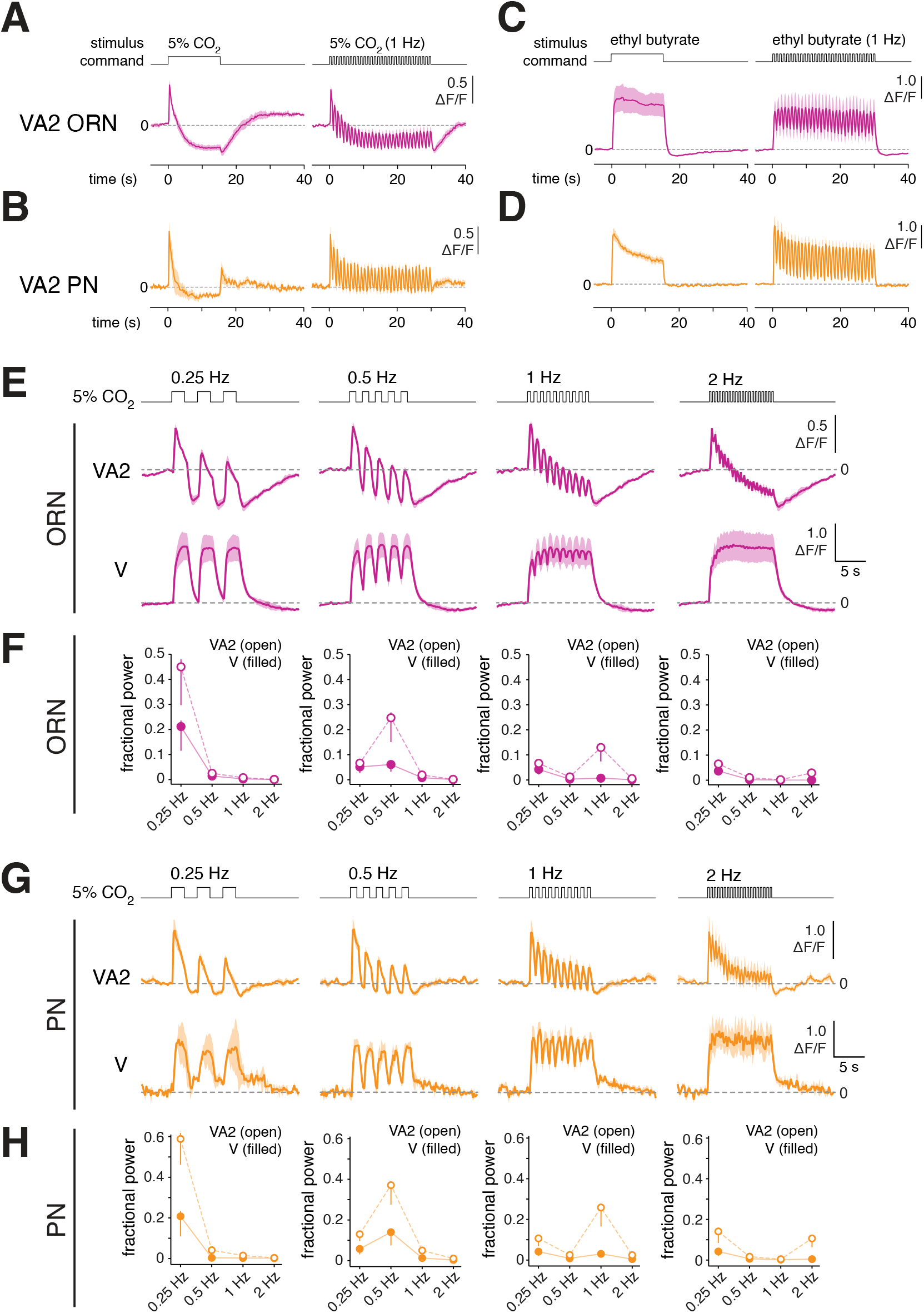
Lateral interactions between ORNs confer odor- and glomerulus-specific response dynamics that are transmitted to downstream PNs. **A-B)** Time course of calcium signals in ORN axon terminals (A, magenta, *n* = 4 flies) or PN dendrites (B, orange, *n* = 5-7 flies) in glomerulus VA2 in response to either a sustained 15 s pulse of 10% CO_2_ (left) or to a 30 s 1 Hz train of 10% CO_2_ pulsed at 50% duty cycle. The top row shows the open-closed state of the odor delivery valve. **C-D)** Same as panels A-B, but in response to the odor ethyl butyrate (10^- 4^). VA2 presynaptic terminals respond phasically to CO_2_ but tonically to ethyl butyrate, and these differences in response dynamics are inherited by downstream PNs. **E)** Time course of calcium signals in ORN presynaptic terminals in glomerulus VA2 or V (*n* = 3 flies) in response to a 10 s train of 10% CO_2_ stimuli presented at 0.25 Hz, 0.5 Hz, 1 Hz, or 2 Hz, pulsed at 50% duty cycle. **F)** Fractional power at each of the indicated frequencies, determined from the power spectral density estimate of the calcium signals immediately above each plot at each stimulus frequency. Error bars are calculated from the 95% confidence bounds of the power spectral density estimate. **G-H)** Same as panels E-F, but for PN dendrites in glomerulus VA2 or V (*n* = 4 for VA2, *n* = 3 for V). GCaMP6f is expressed in either all ORNs (from *pebbled-gal4* for VA2 and V ORNs in panels A, C, E-F), a large subset of PNs (from *GH146*-*gal4* for VA2 PNs in panels B, D, G-H), or in V PNs (from *VT12760*-*gal4* in panels G-H).

Whereas VA2 ORNs responded phasically to long pulses of CO_2_ responses (Figure 6A), we observed that V ORNs responded tonically to CO_2_ (Figure 1F). These observations suggested that VA2 ORNs apply a filter to CO_2_ stimuli that preferentially transmits higher frequency CO_2_ fluctuations to downstream neurons, and V ORNs preferentially transmit slower CO_2_ fluctuations. To directly test this prediction, we delivered pulses of CO_2_ at varying frequencies to flies while measuring calcium signals in VA2 or V ORNs and PNs (Figure 6E, G). At low frequencies, presynaptic calcium in V ORNs closely followed the stimulus, more accurately representing the absolute levels of CO_2_ as compared to VA2 ORNs, which adapted strongly. However, as the stimulus frequency was increased, presynaptic calcium in V ORNs failed to keep pace with the stimulus and encoded stimulus fluctuations with decreasing contrast compared with VA2 ORNs (Figure 6E). At higher stimulus frequencies (> ∼2 Hz), the timescale of the stimulus fluctuations approaches that of the kinetics of the calcium indicator (τ1/2 ∼ 200-400 ms) (Chen et al., 2013), and so the temporal resolution observed for VA2 responses to CO_2_ represents a lower bound. Measurements of calcium signals in VA2 and V PNs to pulses of CO_2_ show that the transient and sustained dynamics of VA2 and V ORN responses, respectively, are transmitted across the synapse (Figure 6G).

We computed the power spectral density estimate of the mean ORN or PN calcium signal in each glomerulus, recorded at each frequency of CO_2_ presentation, and determined how much of the signal power was contained at the stimulus frequency. We found that the fractional signal power at the stimulus frequency was significantly higher for VA2 ORNs or PNs, as compared to their respective signals in glomerulus V, and this effect became more pronounced as the stimulus frequency increased, up to 1 Hz (Figure 6F, H). This analysis shows that VA2 ORNs, and subsequently VA2 PNs, are more effective at encoding fast fluctuations in CO_2_ concentration as compared to the equivalent cells in glomerulus V.

Taken together, these results demonstrate that CO_2_ is encoded in parallel by V ORNs and VA2 ORNs which impose different temporal filters on the stimulus. Calcium signals in slowly adapting V ORNs more accurately reflect the absolute concentration of CO_2_ integrated over relatively long timescales (seconds), whereas rapidly adapting VA2 ORNs preferentially transmit fast changes in CO_2_ levels. These differences in presynaptic stimulus dynamics are inherited by postsynaptic PNs in the brain, which is largely expected since ORN presynaptic calcium is tightly coupled to neurotransmitter release. As a result, the dual representation of CO_2_ in glomeruli with distinct response dynamics expands the coding bandwidth of the circuit for CO_2_ stimuli.

## DISCUSSION

### Functions of stimulus-selective lateral information flow between ORNs

Sensory integration is commonly studied in central circuits, but substantial crosstalk occurs between primary sensory neurons before sensory information enters the brain. In this study, we describe a novel form of lateral information flow, comprised of mixed excitation and inhibition between primary olfactory afferents, which enables stimulus-specific integration of sensory information and markedly reshapes the spatiotemporal structure of odor representations. This finding expands the list of circuits in which lateral excitatory coupling occurs between primary sensory neurons, which includes invertebrate primary mechanosensory neurons, vertebrate photoreceptors, and primate peripheral nociceptive fibers (Bloomfield and Völgyi, 2009; Meyer et al., 1985; Rabinowitch et al., 2013), and suggests that such lateral interactions between primary afferents may be a common feature of early sensory circuits. We demonstrate that ORN presynaptic calcium signals can be driven by two sources of excitation: 1) direct excitation derived from odorant receptor-mediated olfactory transduction, and 2) lateral input derived from specific subsets of other ORNs. Thus, excitation originating from different classes of olfactory receptors is combined and processed before it even leaves the primary afferents, a finding that has implications for traditional “labeled line” theories of chemosensory coding (Galizia, 2014; Haverkamp et al., 2018).

The novel form of lateral signaling between ORNs is distinguished by its selectivity – whereas previously described forms of lateral connectivity in the antennal lobe predominantly connect glomeruli in an all-to-all manner (Hong and Wilson, 2015; Olsen and Wilson, 2008; Olsen et al., 2007; Shang et al., 2007; Silbering and Galizia, 2007; Silbering et al., 2008), this new form of lateral signaling enables subsets of glomeruli to interact in a selective and directional manner and has several important functional consequences for odor coding. First, stimulus-selective lateral input enters ORNs downstream of action potential generation, and, thus, can lead to odor-specific decoupling of levels of synaptic transmission from firing rate in ORNs (Figure 3B, 5B). This fact is most dramatically demonstrated by the presence of strong odor-evoked calcium signals at the presynaptic terminals of silenced ORNs that cannot spike due to mutation of their cognate OR (Figure 5D). This result highlights the role of individual ORNs as complex, nonlinear computational units which integrate and profoundly transform olfactory input through both odorant interactions at the OR (Pfister et al., 2020; Xu et al., 2020) and through lateral signaling (see also (Su et al., 2012).

Second, stimulus-selective lateral signaling contributes to the broadening of PN odor tuning compared to its cognate presynaptic ORN (Bhandawat et al., 2007; Wilson et al., 2004). PN broadening has been attributed to nonlinearities in the intraglomerular ORN-to-PN transformation, driven by signal amplification from convergent ORN input (Bhandawat et al., 2007), synaptic depression at ORN-PN synapses (Kazama and Wilson, 2008), and global lateral inhibition that is mediated by local GABAergic neurons (Olsen and Wilson, 2008; Olsen et al., 2010; Root et al., 2008; Silbering and Galizia, 2007; Silbering et al., 2008). In addition, dense electrical coupling between PNs may boost PN responses to weak inputs (Olsen et al., 2007; Shang et al., 2007; Yaksi and Wilson, 2010). These mechanisms act mostly uniformly and globally across all glomeruli to scale glomerular output according to the current levels of activity in the network. In contrast, glomerulus-selective lateral signaling leads to odor-specific broadening only within specific subsets of glomeruli. These glomeruli may be related by the functional relationships between the odors they directly detect, and thus selective lateral inputs may implement broadening that supports the behavioral logic of the interacting chemical cues (see below). Selective lateral interactions such as these may contribute to previously observed differences in the rank order of odor preferences between cognate ORNs and PNs in the same glomerulus (Bhandawat et al., 2007), which are difficult to account for by globally acting mechanisms.

Third, the new form of stimulus-specific lateral signaling enables previously undescribed synaptic computations in the olfactory circuit. The interplay of lateral input with direct excitation modifies the dynamics of odor-driven calcium signals in a manner that can depend on odor or glomerulus identity. For instance, in glomerulus VA2, the distinct response dynamics elicited by fluctuating ethyl butyrate versus CO_2_ demonstrate that glomerular output dynamics can encode chemical identity information (Figure 6A-D), a general principle that has been suggested in other olfactory systems (Laurent, 1999; Uchida et al., 2014). Furthermore, lateral information flow can generate parallel versions of a stimulus in multiple glomeruli, but which are differentially filtered compared to the original (Figure 6E). The different stimulus dynamics that are preferentially captured by each glomerulus may be matched to the distinct olfactory behaviors to which each is coupled (see below). In this way, stimulus-selective lateral information flow expands the coding bandwidth of the circuit, capturing stimulus fluctuations across a wider range of timescales relevant to different odor-driven behavioral responses.

### Potential mechanisms of lateral signaling between ORNs

Stimulus-selective lateral signaling between ORNs does not appear to require any of the major classes of chemical neurotransmitters in the antennal lobe (Figure S4A-B), nor does it require the function of the innexin *shakB*, which has been previously implicated in electrical coupling in the antennal lobe (Yaksi and Wilson, 2010). However, lateral signaling could be mediated by electrical synapses comprised of any of the other seven innexins in *Drosophila* (Stebbings et al., 2002). Another possibility is that lateral ORN interactions could be mediated by electric field effects between adjacent ORN axons arranged in an electrically isolated extracellular space (Faber and Pereda, 2018). This phenomenon, known as ephaptic coupling, occurs bidirectionally, but asymmetrically between ORN cell bodies grouped in the same antennal sensillum, and mediates lateral inhibition of ORN spiking (Su et al., 2012; Zhang et al., 2019). In the mammalian olfactory system, biophysical modeling predicts that ephaptic coupling between olfactory sensory axons would allow an action potential in one axon to elicit action potentials in another within the olfactory nerve (Bokil et al., 2001). Notably, CO_2_ activation of ab1C ORNs elicits excitation in all three of the other glomeruli (DM1, VA2, and DL1) targeted by ORN classes housed in the ab1 sensillum (ab1A, ab1B, and ab1D, respectively). The axons of ORN classes grouped together in a common sensillum may fasciculate together as they project through the antennal nerve, facilitating local field effects. However, additional glomeruli that are not targeted by ab1 ORN classes are also excited by ab1C activation (Figure 1A, and data not shown), and observations from the DM1 glomerulus (Figure 5B, D) suggest that ORNs need not be grouped in a sensillum for lateral excitation to exist between them. Understanding the physical arrangement among axons belonging to different classes of ORNs in the antennal nerve may provide insight into the logic of their lateral coupling. Lastly, we did observe that the excitatory component of lateral input is sensitive to ouabain (Figure S4D). Inhibition of the Na^+^/K^+^-ATPase pump has many possible effects, which could include the loss of the electrogenic current, disruption of the unusual ionic gradients in insect sensilla (Kaissling and Thorson, 1980), and/or an increase in intracellular calcium (Tian and Xie, 2008). More work will be required to understand which of these contributes to a loss of signaling between ORNs.

Another possible mechanism that could mediate lateral signaling between ORNs is peptidergic or neuromodulatory signaling. Prior work shows that presynaptic ORN calcium can be modulated by neuropeptides such as short neuropeptide F and tachykinin, depending on satiety state (Ko et al., 2015; Root et al., 2011; Su and Wang, 2014). In addition, the antennal lobe is densely innervated by local neurons, most of which project globally to all glomeruli, but some of which innervate only limited subsets of glomeruli (Chou et al., 2010; Wilson, 2013); the latter could serve as substrates for selective interactions between glomeruli. Serial electron microscopy reconstruction of antennal lobe circuits shows that the majority of local neuron profiles have synaptic release sites containing dark dense core vesicles (Horne et al., 2018), which typically carry neuropeptides, monoamines, or other modulatory molecules. Thus, many local neurons likely release additional transmitters or modulators in addition to GABA. Further analysis of the full antennal lobe connectome may reveal modulatory neurons that selectively innervate the subset of glomeruli connected by these lateral interactions and that would therefore be candidate intermediary substrates.

The hypothesis that a neuromodulatory neuron might mediate lateral ORN signaling is in part motivated by the observation that CO_2_-driven lateral signals are modulated from a low to a high responsive state during the experiment. We cannot rule out that the low state is a byproduct of exposing the brain, which is unavoidable for experimental recordings; however, we emphasize that conventional odor responses (arising from direct OR activation and spiking) are present and unchanged in amplitude or dynamics from the start of the experiment to its conclusion. One possibility is that potentiation of the laterally evoked responses reflects a change in the animals’ internal state, for instance, adaptation to the initial stress caused by preparing the animal for recordings. We attempted a number of coarse manipulations of the flies’ internal state prior to recording, including starvation, sucrose feeding, exposure to extreme temperatures, recording at different timepoints of the circadian cycle, and social crowding. However, with our sample sizes (∼20 flies), we were unable to identify a manipulation that consistently biased the flies into either the low or high state (data not shown).

The lability of CO_2_-evoked ORN activity is notable in the context of the state-dependent effects of CO_2_ on fly behavior (Bräcker et al., 2013). Flies behave differently in response to CO_2_ depending on their satiety state, circadian state, temperature, and walking speed. For instance, ten minutes after they are first introduced into an enclosed walking arena, flies avoid a local source of CO_2_; however, if allowed to acclimate in the arena for two hours, flies exhibited attraction to CO_2_ (van Breugel et al., 2018). Further work aimed towards linking the modulation of CO_2_ responses in ORN terminals with the animals’ ongoing behavioral state will help to more precisely elucidate their relationship.

### The role of lateral ORN signaling in the processing of CO_2_ cues

CO_2_ is a complex environmental cue for flies that can elicit behavioral aversion in some contexts (Ai et al., 2010; Faucher et al., 2006; Jones et al., 2007; Lin et al., 2013; Suh et al., 2004) and attraction in others (van Breugel et al., 2018; Wasserman et al., 2013). Whereas aversive responses to CO_2_ are known to be mediated by signaling through Gr63a/Gr21a and the V glomerulus (Jones et al., 2007; Kwon et al., 2007; Suh et al., 2004), the odorant receptors that mediate attraction to CO_2_ are still unknown. CO_2_-driven lateral excitation in ORNs in VA2 and DM1 is notable given that these glomeruli are activated by, and mediate attraction to, food odors like apple cider vinegar (de Bruyne et al., 2001; Faucher et al., 2013; Semmelhack and Wang, 2009). However, CO_2_- evoked lateral excitation in VA2 and DM1 cannot be solely responsible for the previously described attraction of flies to CO_2_ (van Breugel et al., 2018), the latter of which depends on neither Gr63a/Gr21a- nor Orco-signaling (see also Figure S4E). We note, however, that the loss of the inhibitory component of CO_2_-evoked lateral signals in *orco^-/-^* flies implies the existence of an Orco-dependent OR that is sensitive to CO_2_ and is the source of the inhibitory component of the lateral response. In fact, a prior study reported that CO_2_ tracking during flight requires the function of Orco (Wasserman et al., 2013). These results highlight the complexity of CO_2_ encoding in the fly brain. The behavioral response to CO_2_ is likely determined by the context-dependent integration of multiple olfactory signals, which include aversion-promoting activity in V, attraction-promoting activity in VA2 and DM1, as well as activity in as-yet-unidentified Ir25a-dependent (van Breugel et al., 2018) and Orco-dependent ORN classes.

What is the functional benefit of broadening CO_2_ representations in the antennal lobe via stimulus-selective lateral signaling between ORNs? In the natural world, CO_2_ is almost always encountered in combination with other odorants. For instance, over-ripe fruit, an important food source for flies, emits a complex mixture of volatiles that includes ethyl acetate, ethanol, acetic acid, diacetyl, and CO_2_ (Faucher et al., 2006; Turner and Ray, 2009; Zhu et al., 2003). Such low molecular weight esters, alcohols, and short chain fatty acids excite glomeruli that mediate behavioral attraction, including DM1 and VA2 (de Bruyne et al., 2001; Faucher et al., 2013; Semmelhack and Wang, 2009), and CO_2_ levels vary depending on the fruit substrate and the stage of ripening and fermentation (van Breugel et al., 2018; Faucher et al., 2006). CO_2_-driven lateral input to DM1 and VA2 may provide additional information about the appropriateness or likely value of the emitting source, increasing the sensitivity of these glomeruli to high value food odors and more robustly activating downstream circuits coupled to attraction.

Finally, our results also show that the interplay of CO_2_-evoked lateral excitation and inhibition in VA2 and DM1 ORNs leads to more transient CO_2_ response dynamics in VA2 and DM1 as compared to the tonic CO_2_ responses in V (Figure 6). The tonic calcium response dynamics elicited by CO_2_ in ab1C presynaptic terminals in V contrast with prior studies that have emphasized how the spiking responses of CO_2_-sensitive insect ORNs adapt very strongly (Faucher et al., 2013; Grant et al., 1995). Whereas ab1C firing rate better encodes the rate of change in CO_2_ concentration (Faucher et al., 2013), ab1C presynaptic calcium tracks the absolute levels of CO_2_ (Figure 6E). A recent study observed a similar behavior in other classes of ORNs and with other odors, suggesting it may be a general feature of the transformation of ORN spike rate to presynaptic calcium (Martelli and Fiala, 2019).

The dual encoding of CO_2_ with distinct dynamics in glomerulus VA2 (and DM1) as compared to glomerulus V could serve two important functions. First, when flies encounter slow, sustained increases in CO_2_ concentration, phasic activity in VA2 and DM1 ensures that the prolonged pulse of CO_2_ is signaled predominantly via the V glomerulus, so that behavioral aversion should dominate. Such slow, sustained CO_2_ cues might be encountered during walking, from close range interactions with exhalations from large animals, overcrowded populations of flies or other arthropods, or potentially dangerous enclosed natural sources of CO_2_, such as a seeping hive or rotting log. This mechanism may serve a complementary function to the inhibitory mechanisms that suppress CO_2_ activation of V ORNs when food odors are present, which have been proposed to enable flies to approach important nutritive sources like fermenting fruit (Su and Wang, 2014). Two such inhibitory mechanisms have been described, the direct antagonism of Gr63a/Gr21a receptor activity by food odors like diacetyl (Turner and Ray, 2009) and ephaptic inhibition of ab1C spiking by activation of ab1A and ab1B ORNs, which are sensitive to food odors (Su et al., 2012).

Second, when CO_2_ is detected as part of a natural odor mixture emanating from a distant source like fermenting fruit, phasic activation of DM1 and VA2 may enhance the ability of flies to respond to fast odor fluctuations in flight, such as those that occur when navigating filaments of odor in a plume at long distances from the odor source (van Breugel and Dickinson, 2014). We note that, in the case of mosquitoes navigating in a wind tunnel, sustained upwind displacement is observed only when mosquitoes are presented with pulsed carbon dioxide; they fail to fly significantly upwind when the airstream is uniformly permeated with a steady concentration of carbon dioxide (Geier et al., 1999; Gillies, 1980). This result shows that the dynamics of CO_2_ stimuli are significant for odor-guided behavior. The parallel representation of CO_2_ in multiple glomeruli, each capturing distinct stimulus dynamics, would enable downstream circuits to select the most appropriate behavioral response to CO_2_ under the wide range of timescales and contexts in which CO_2_ is encountered in the natural environment.

### The roles of lateral interactions in olfactory processing

Although this study largely focused on the lateral flow of olfactory signals from CO_2_-detecting ab1C ORNs, our results suggest that selective lateral interactions between ORNs may be a general feature of the antennal lobe circuit. Multiple, but not all, odors can elicit activity in the terminals of DM1 ORNs lacking a functional odorant receptor (Figure 5). These odors are diverse and activate a broad range of different glomeruli in the antennal lobe to varying extents, implying the existence of privileged, directional interactions between multiple subnetworks of glomeruli. An interesting question is the relative significance of lateral information flow for reshaping the representation of all odors in the antennal lobe versus specific specialist odors that narrowly activate only one or a few ORs. Despite its narrow input into the circuit, CO_2_ elicits robust population activity in third-order olfactory areas like the mushroom body (Bräcker et al., 2013) at response rates comparable to those of generalist odors activating multiple ORs (∼5%, data not shown). Because third-order mushroom body neurons normally require coincident activity from multiple glomeruli to achieve firing threshold (Gruntman and Turner, 2013), the lateral spread of olfactory input from a glomerulus to a subset of other glomeruli may be especially important in enabling narrowly activating odors to efficiently elicit third-order representations.

This study is one of the most direct demonstrations of selective lateral interactions between specific glomeruli in an olfactory circuit. The highly ordered organization of the antennal lobe, combined with genetic access to identified cell types, allowed the mapping of the identities and odor selectivity of a subnetwork of preferentially connected glomeruli. In the vertebrate olfactory bulb, evidence for selective lateral interactions is mixed (Aungst et al., 2003; Economo et al., 2016; Fantana et al., 2008; Laurent, 1999; Luo and Katz, 2001) and has focused on selective inhibition between glomeruli, indirectly inferred from anticorrelated neural activity. Recent studies suggest that inhibitory interactions between olfactory bulb glomeruli are sparse and selective, linking glomeruli tuned to odors that do not share obvious functionally related properties (Economo et al., 2016; Fantana et al., 2008; Zavitz et al., 2020). This result argues against a conventional role for sparse inhibition in mediating contrast enhancement in the olfactory bulb but leaves unanswered what functions it may serve. In the antennal lobe, selective lateral interactions are occurring via a different circuit mechanism, because they are non-GABAergic and are comprised of mixed excitation and inhibition with varying dynamics. However, this study shows how, in the case of CO_2_, selective lateral signaling can nonlinearly reformat sensory input to the antennal lobe in ways that can support the chemical ecological functions of this important environmental cue. It may provide a useful guide for future investigations into the broader functions of selective lateral interactions in olfactory circuits.

## AUTHOR CONTRIBUTIONS

**Dhruv Zocchi:** Conceptualization, Methodology, Investigation, Validation, Formal analysis, Writing – Original Draft, Writing – Review & Editing, Visualization. **Emily S. Ye:** Methodology, Investigation, Validation, Formal analysis. **Elizabeth J. Hong:** Conceptualization, Methodology, Formal analysis, Writing – Original Draft, Writing – Review & Editing, Visualization, Supervision, Funding acquisition.

## ACKNOWLEDGMENTS

We are grateful to B. D. Pfeiffer and D. J. Anderson for gifts of unpublished fly stocks. We thank T. Lee, and R. J. Wyman for sharing fly stocks, and we thank G. Rubin and L. M. Stevens for sharing plasmids. We thank A. Dea for the generation of the *lexAop-DTI* flies. We thank M. Meister and P. W. Sternberg for helpful comments, and M. H. Dickinson and members of the Hong lab for careful readings of the manuscript. We especially acknowledge F. van Breugel and M. H. Dickinson for stimulating conversations and for sharing unpublished results which partially motivated this study. This work was funded by grants to E. J. H. from the National Science Foundation (Ideas Lab 1556230), the National Institutes of Health (1U01MH109147), and the Shurl and Kay Curci Foundation. E. J. H. is a Clare Boothe Luce Professor of the Henry Luce Foundation.

## SUPPLEMENTAL FIGURE LEGENDS

**Figure S1:**
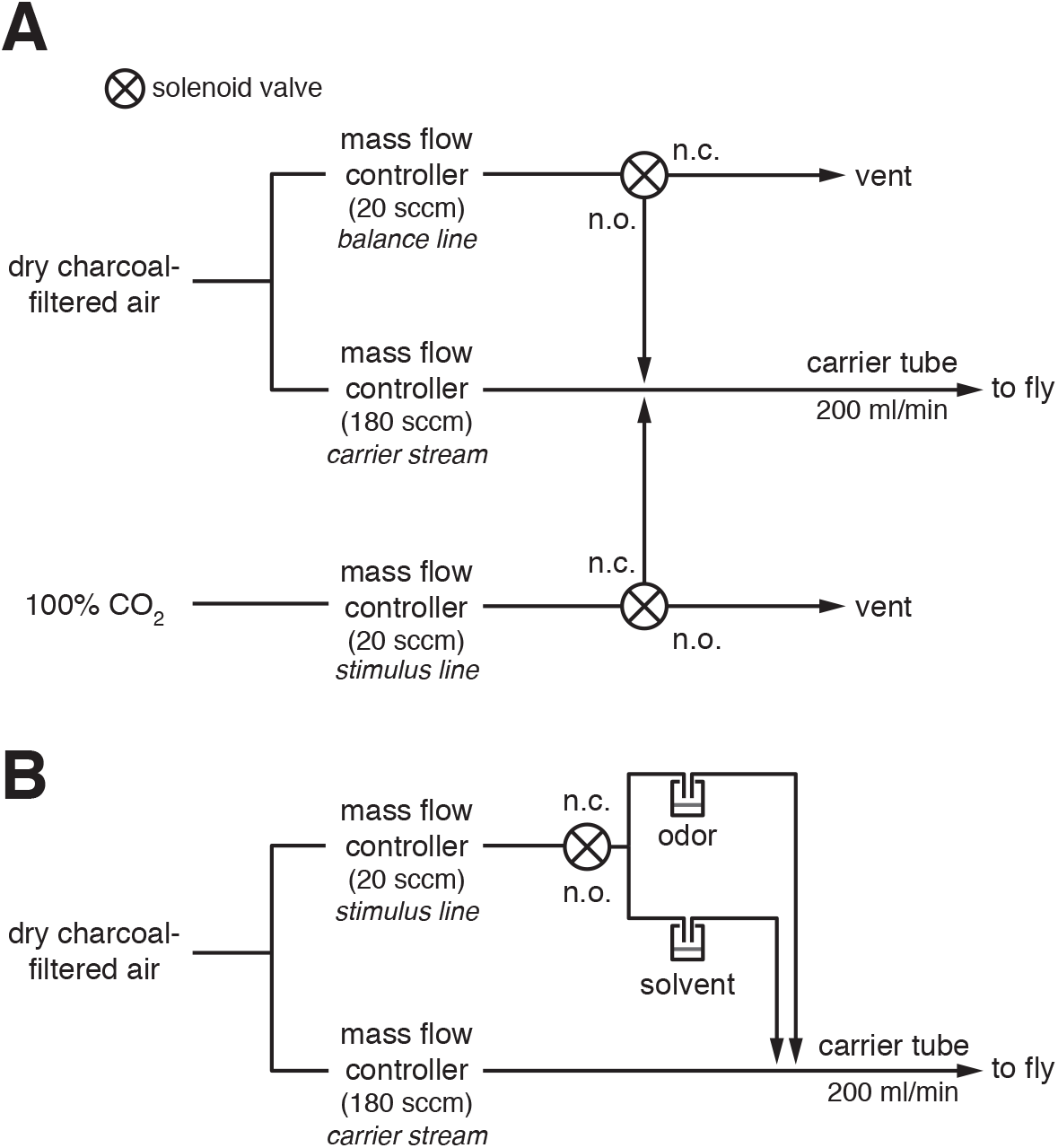
Odor delivery (related to Figure 1). **A)** Olfactometer design for delivery of 10% CO_2_ stimulus. The flow rates of the stimulus and balance lines were always kept equal. The normally open (n.o.) outlet of each solenoid valve is open when the valve is unpowered and is closed when the valve is powered. The normally closed (n.c.) outlet of each solenoid valve is closed when the valve is unpowered and is open when the valve is powered. The same command signal is provided simultaneously to both valves. CO_2_ stimuli of varying concentration were generated by adjusting the relative flow rates of the carrier line, and the stimulus and balance lines. For instance, for a 5% CO_2_ stimulus, the mass flow controller for the carrier line was set to 190 sccm, and the stimulus and balance lines were set to 10 sccm. A total flow rate of 200 ml/min at the fly was used for all experiments, except for the optogenetic experiments in Figure 4E-G and the experiments in Figure 6, where a total flow rate of 2 L/min was used. **B)** Olfactometer design for delivery of all other odor stimuli besides CO_2_ (see Method Details). Odor concentrations in the figures and text are reported as the v/v dilution of the odor in solvent in the vial. Odors were diluted 10-fold in air compared to their concentration in the headspace of the vial. The only exceptions to this were acetic acid, which was always diluted 2-fold, and ammonia, which was diluted 100-fold in Figure 1, in air. Diagram is not drawn to scale.

**Figure S2:**
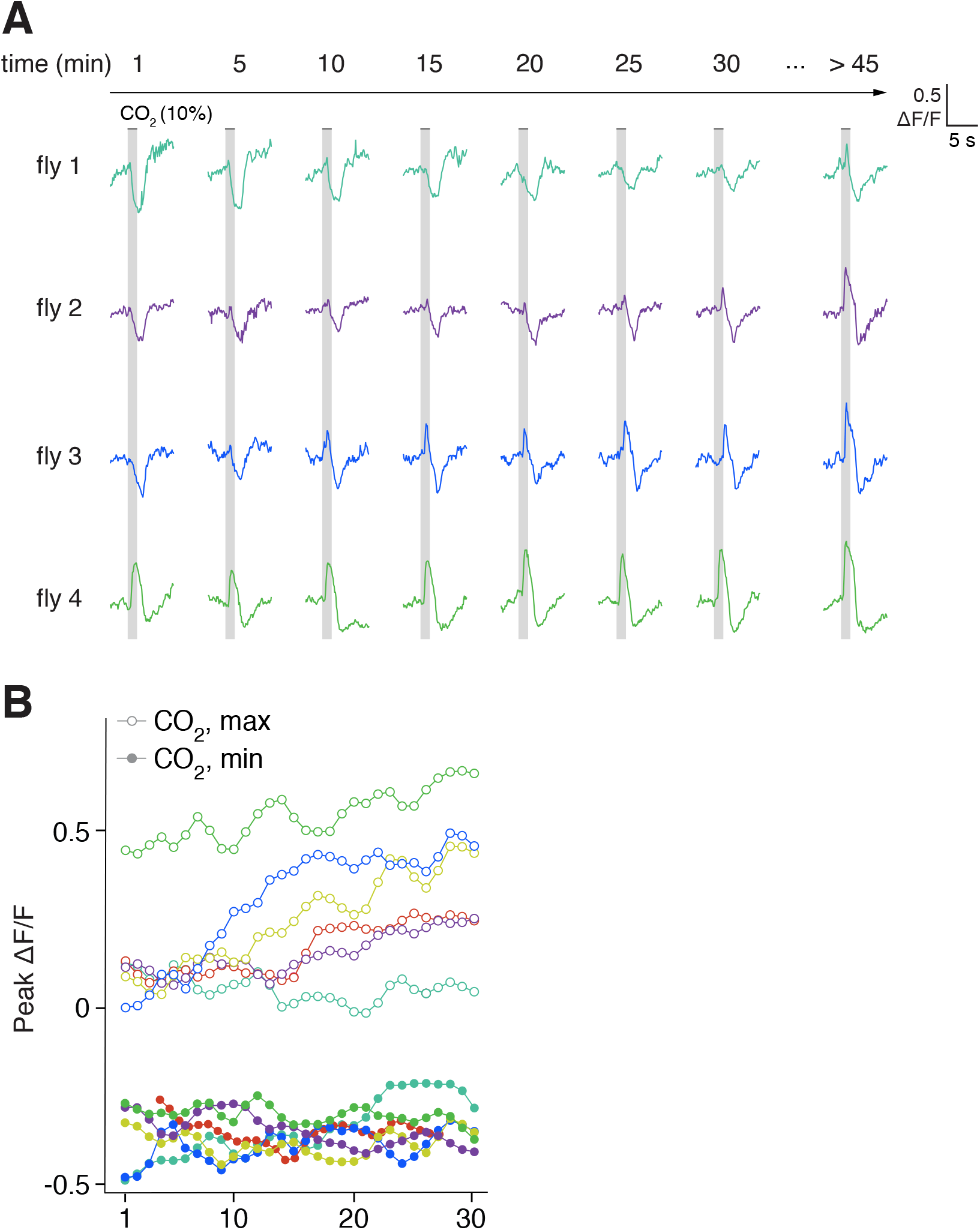
Variation in the time course of modulation of CO_2_ responses in ORN terminals in VA2 from a low to high state (related to Figure 2). **A)** Single trial recordings of the change in fluorescence in ORN terminals in VA2 elicited by CO_2_ over the duration of an experiment in four individual flies. A 3 s pulse of 10% CO_2_ was presented once every minute for 30 presentations; every fifth trial is shown. The fly was then left unstimulated on the rig, and a final measurement was taken between 45-60 min from the start of the experiment. **B)** Peak amplitudes of single-trial responses to CO_2_ in ORN terminals in VA2, measured as in A, for six representative flies (which include the four in A). Both maximum peaks (open symbol) and minimum peaks (closed symbol) are shown. Note the variation in the time course of modulation of the CO_2_ response across different experiments. During the 30-minute recording period, the recording began with responses to CO_2_ in ORN terminals in VA2 already in the “high state” for one fly (fly 4, green), the response was mostly unchanged for one fly (fly 1, teal), and the CO_2_ responses in the remaining four flies were significantly modulated over the course of the recordings, though at varying rates.

**Figure S3:**
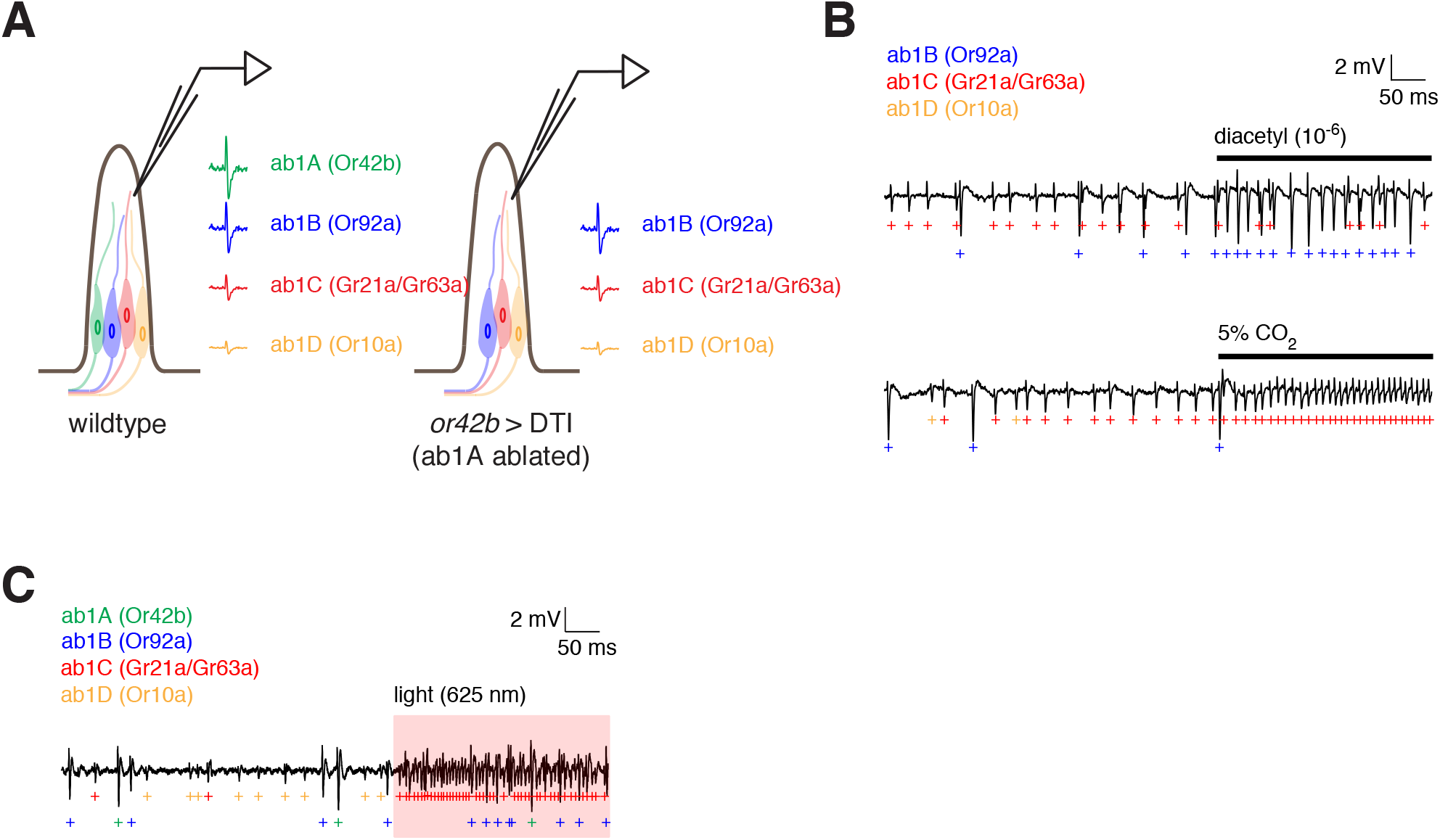
CO_2_ elicits spiking responses in ab1C, but not ab1B, ORNs (related to Figures 3 and 4). **A)** Schematic of experimental setup. The wildtype ab1 sensillum (left) houses four classes of ORNs – ab1A, ab1B, ab1C, and ab1D – which have characteristic spike amplitudes of descending size in extracellular single sensillum recordings (SSR). The two largest spikes, originating from the ab1A and ab1B, have the most similar amplitudes and are sometimes challenging to accurately assign to their respective ORN class, whereas spikes for the remaining two ORN classes, ab1C and ab1D, are easily identifiable and reliably sorted to their respective ORN class. To unambiguously identify ab1B spikes, we genetically ablated ab1A ORNs by expressing diphtheria toxin under the control of the *or42b-gal4* driver (right). When ab1A was absent, ab1B spikes were easily differentiated from ab1C and ab1D spikes based on shape and amplitude. Note that, in this fly, GCaMP6f was also expressed in most ORNs under the control of *orco-gal4* to allow imaging of calcium signals in ORN terminals in the same fly (Figure 3). **B)** Example SSR recordings from the ab1 sensillum in flies in which ab1A ORNs were ablated (panel **A**, right), showing the spiking responses of ab1B ORNs to 10^-6^ diacetyl (top) and the spiking responses of ab1C ORNs to 5% CO_2_ (bottom). Note that no ab1B spikes (blue) were evoked when flies were presented with CO_2_. **C)** Example SSR recording from the ab1 sensillum in flies expressing CsChrimson under the control of *gr21a-gal4* and GCaMP6f under the control of *orco-gal4* (see Figure 4E). Exposing the antenna to red light (625 nm, red shading) elicits spiking selectively in ab1C ORNs. “+” symbols mark identified spikes from each ORN class in the ab1 sensillum.

**Figure S4:**
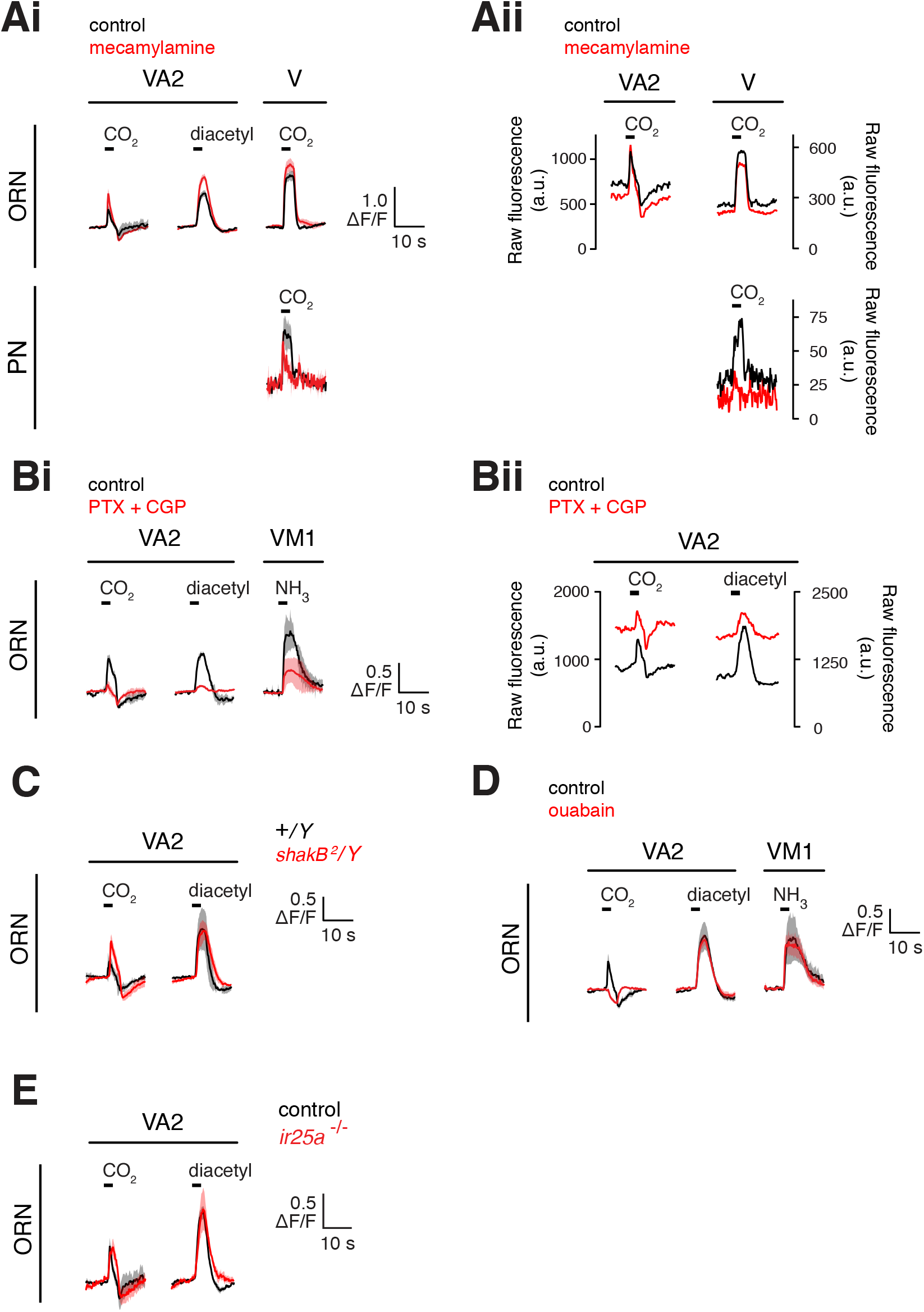
Pharmacological and genetic investigation of the mechanism of lateral signal flow between ORNs (related to Figure 4). **A)** Odor-evoked calcium signals in either ORN terminals (top) or PN dendrites (bottom) in glomeruli VA2 or V before (black) and after (red) bath application of the cholinergic antagonist mecamylamine (100 μM). The *Δ*F/F response across flies (*n*=4, mean and s.e.m.) is shown in **Ai**, and a representative example of the raw odor-evoked fluorescence signal from one experiment is shown in **Aii**. Although mecamylamine reduced CO_2_-evoked responses in postsynaptic PN dendrites in glomerulus, as expected, both the excitatory and inhibitory components of the CO_2_ response in ORN terminals in VA2 were largely unaffected. The small enhancement of some odor responses in VA2 ORNs in mecamylamine could be secondary to a loss of presynaptic inhibition onto ORNs due to blockade of ORN to GABAergic local neuron signaling. **B)** Odor-evoked calcium signals in ORN terminals in glomeruli VA2 or VM1 before (black) and after (red) bath application of the GABAAR and GABABR antagonists picrotoxin (5 μM) and CGP54626 (50 μM), respectively. The *Δ*F/F response across flies (*n*=2, mean and s.e.m.) is shown in **Bi**, and an example of the raw odor-evoked fluorescence signal from one experiment is shown in **Bii**. Wash-in of GABA receptor antagonists led to a large increase in the baseline fluorescence in ORN terminals, likely reflecting the loss of tonic presynaptic inhibition. Although this led to a reduction in the dynamic range of the calcium indicator and a diminishment of the size of all odor responses, both the excitatory and inhibitory components of the CO_2_ response in ORN terminals in VA2 clearly persisted in the presence of GABA receptor blockers. **C)** Odor-evoked responses (mean and s.e.m.) to CO_2_ or diacetyl in ORN terminals in glomerulus VA2 in control (black, n=2) or *shakB^-/y^* mutant (red, *n*=4) flies. The CO_2_ response in VA2 ORNs does not appear to require *shakB* function. **D)** Odor-evoked calcium signals (mean and s.e.m., *n*=4 flies) in ORN terminals in glomeruli VA2 or VM1 before (black) and after (red) bath application of the Na^+^/K^+^-ATPase inhibitor ouabain (100 μM). Inhibition of the Na^+^/K^+^-pump resulted in the selective loss of the excitatory component of CO_2_ responses in ORN axon terminals in VA2, although the inhibitory component of these responses persisted. In addition, ORN responses to other odors (diacetyl in VA2 and ammonia in VM1) that do not depend on lateral signaling between ORNs were unaffected by application of ouabain. **E)** CO_2_- or diacetyl-evoked responses (mean and s.e.m.) in ORNs in glomerulus VA2 in control (black, n=4) or *ir25a*^-/-^ mutant (red, *n*=3) flies. The CO_2_-evoked response in VA2 ORNs does not depend on *ir25a* function. Behavioral genetic experiments indicate that CO_2_ attraction depends on the activity of *ir25a* (van Breugel et al., 2018), which encodes a co-receptor for a subset of the ionotropic class of olfactory receptors (Benton et al., 2009; Silbering et al., 2011). Neither V nor VA2 nor DM1 ORNs are Ir25a-dependent (Benton et al., 2009; Silbering et al., 2011), and CO_2_-evoked lateral signals in ORNs in DM1 (data not shown) and VA2 are present in *ir25a*^-/-^ flies. These results are inconsistent with a role for CO_2_-evoked lateral signaling in VA2 and DM1 ORNs in mediating the behavioral attraction to CO_2_ described in (van Breugel et al., 2018).

**Figure S5:**
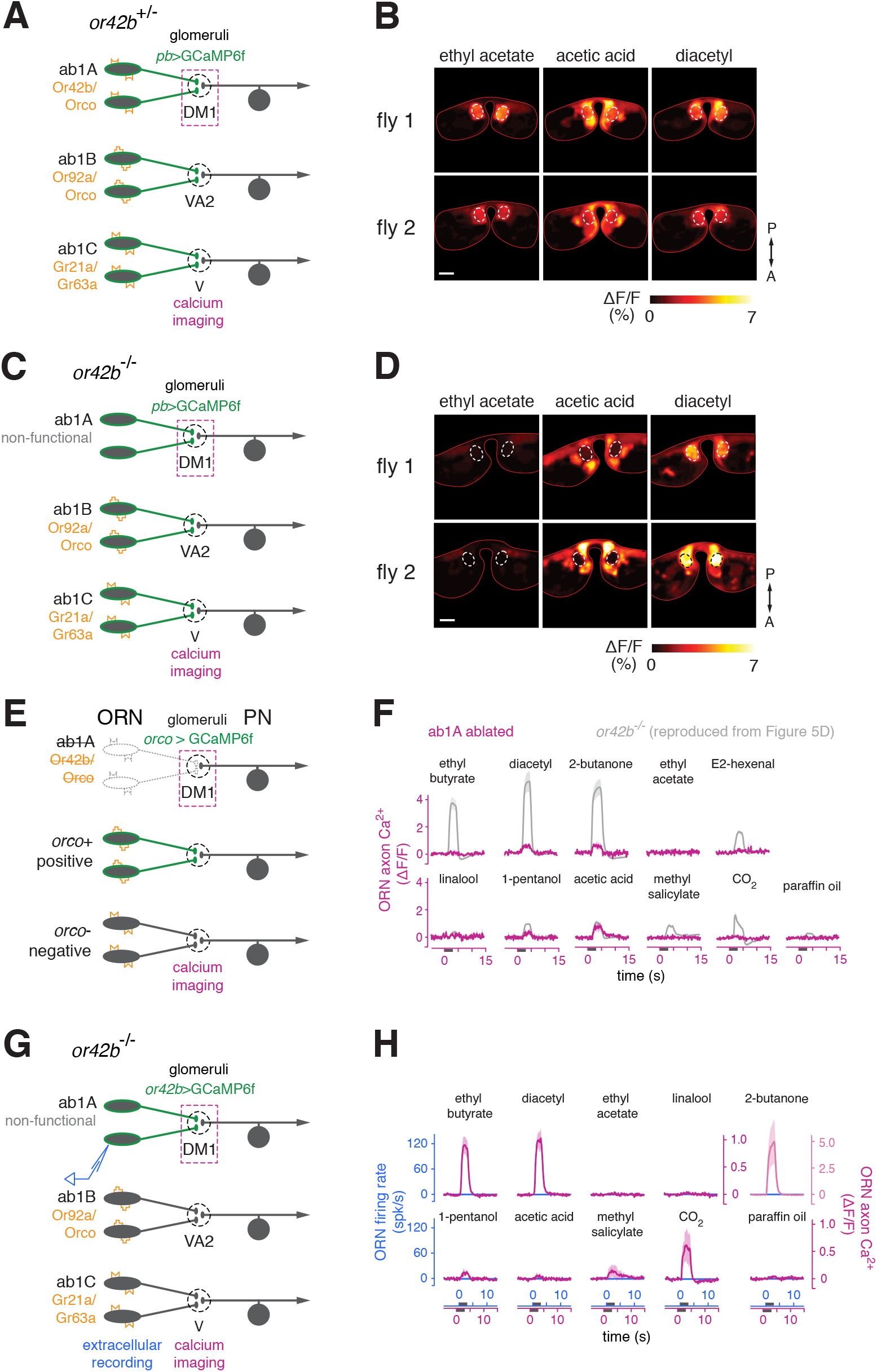
Odors evoke calcium signals in the axon terminals of functionally silent ab1A ORNs lacking odorant receptors (related to Figure 5). **A)** Schematic of experimental setup. Control recordings in *or42b^+/-^* heterozygous flies expressing OpGCaMP6f in all ORNs from the *pebbled-gal4* driver. **B)** Example peak *Δ*F/F responses in ORN terminals from two individual *or42b^+/-^* flies to ethyl acetate (10^-8^), acetic acid (3%), and diacetyl (10^-4^) in the imaging plane containing glomerulus DM1 (dashed white line). Note that all three odors evoke responses in ORN terminals in DM1 in the control genotype. Scale bar is 20µm. **C)** Schematic of experimental setup. Recordings were performed in *or42b^-/-^* flies expressing GCaMP6f in all ORNs (from *pebbled-gal4*). These flies lack a functional odorant receptor in ab1A neurons (which target DM1), thereby abolishing all olfactory transduction and odor-evoked spiking in these ORNs (Figure 5D, blue traces). **D)** Same as B, but in two example *or42b^-/-^* flies. Note the markedly reduced responses of ORN terminals in DM1 to ethyl acetate (10^-8^) and acetic acid (3%), but the strong response to diacetyl (10^-4^) (see also Figure 5D). The odor stimuli were presented at the same concentrations as in Figure 5. **E)** Schematic of experimental setup. ab1A ORNs were ablated by expressing diphtheria toxin under the control of the *or42b-gal4* driver. Flies also expressed GCaMP6f in most ORNs under the control of *orco-gal4*, including in ab1A ORNs if present. **F)** Time course of changes in fluorescence (*Δ*F/F, mean and s.e.m.) in glomerulus DM1 in response to a panel of odorants (same as in Figure 5), recorded in the same imaging plane as in Figure 5B, D, F. The analogous responses from *or42b*^-/-^ flies are overlaid in grey (replotted from Figure 5D) for reference; the comparative absence of responses when ab1A was ablated (magenta, *n*=4) demonstrates that responses from the presumed region of interest containing DM1 in this imaging plane are indeed stemming from ab1A ORNs. **G)** Schematic of experimental setup. The *or42b*^-/-^ mutation was introduced into flies expressing GCaMP6f selectively in ab1A ORNs under the control of *or42b-gal4*, allowing unambiguous identification of ab1A axon terminals in a fly where the ab1A ORN lacks an odorant receptor and is functionally silent. **H)** Comparison of odor-evoked ab1A spiking responses (blue) and the time course of odor-evoked change in fluorescence (*Δ*F/F, mean and s.e.m.) in ab1A axon terminals (magenta, *n* = 3). In this fly, ab1A ORNs did not spike in response to odor (blue PSTHs) but ab1A ORN terminals responded robustly to many odors. This result shows that many odors can recruit lateral input to ab1A ORN terminals.

**Figure S6.**
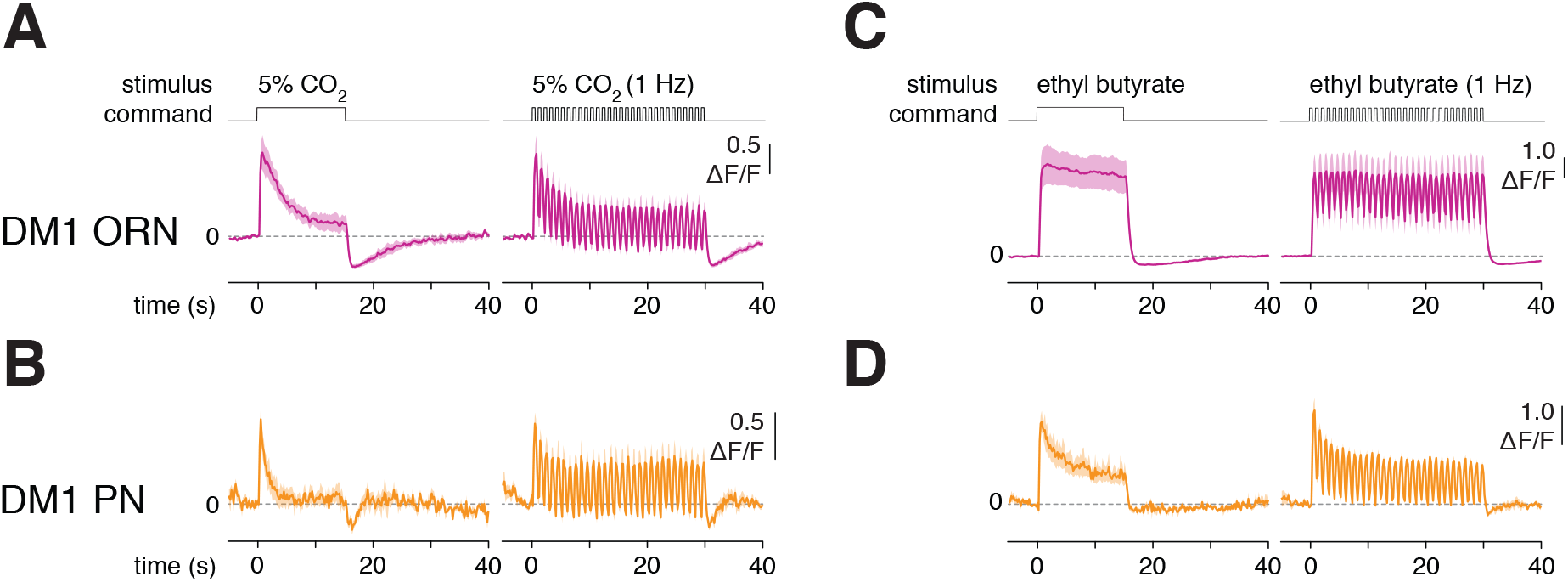
Lateral information flow between ORNs confers odor-specific response dynamics in glomerulus DM1 that are transmitted to downstream PNs (related to Figure 6). **A-B)** Time course of calcium signals in ORN axon terminals (A, magenta, *n* = 4 flies) or PN dendrites (B, orange, *n* = 5-7 flies) in glomerulus DM1 in response to either a sustained 15 s pulse of 10% CO_2_ (left) or to a 30 s 1 Hz train of 10% CO_2_ pulsed at 50% duty cycle. The top row shows the open-closed state of the odor delivery valve. **C-D)** Same as panels A-B, but in response to the odor ethyl butyrate (10^-4^). Calcium signals in DM1 ORN presynaptic terminals adapted rapidly in response to CO_2_, whereas the signal was more sustained in response to ethyl butyrate. These differences in ORN response dynamics were inherited by downstream PNs.

## STAR METHODS

### Resource Availability

#### Lead Contact

Further information and requests for resources should be directed to and will be fulfilled by the Lead Contact, Elizabeth J. Hong

#### Materials Availability

This study generated a new transgenic line, *lexAop-DTI* (III). This stock will be deposited in the Bloomington *Drosophila* Stock Center collection.

#### Data and Code Availability

All data and analysis scripts will be made available to any researcher for the purposes of reproducing or advancing the results.

### Experimental Model and Subject Details

*Drosophila melanogaster* were raised on a 12:12 light:dark cycle at 25°C and 70% relative humidity on cornmeal/molasses food containing: water (17.8 l), agar (136 g), cornmeal (1335.4 g), yeast (540 g), sucrose (320 g), molasses (1.64 l), CaCl2 (12.5 g), sodium tartrate (150 g), tegosept (18.45 g), 95% ethanol (153.3 ml) and propionic acid (91.5 ml). All experiments were performed in female flies 5-20 days post-eclosion, with the exception of experiments in the *shakB^2^* genotype, which were performed in hemizygous males. The specific genotypes of the flies used for each figure panel are given in Table S1.

### Method Details

#### Fly stocks

The transgenes used in this study were acquired from the Bloomington Drosophila Stock Center (BDSC), unless otherwise noted, and have been previously characterized as follows: *pebbled-gal4* expresses Gal4 in all ORNs (Sweeney et al., 2007); *orco-lexA* was from T. Lee and expresses LexA under the control of the *orco* promoter (Lai and Lee, 2006); *or42b-gal4* expresses Gal4 in ab1A ORNs (Fishilevich and Vosshall, 2005); *or92a-gal4* expresses Gal4 in ab1B ORNs (Fishilevich and Vosshall, 2005); *gr21-gal4* expresses Gal4 in ab1C ORNs (Couto et al., 2005); *GH146-gal4* expresses Gal4 in a large subset of PNs, including DM1 and VA2 PNs (Stocker et al., 1997); *VT12760*-*gal4* (III) expresses Gal4 in multiglomerular PNs with dendrites in glomerulus V (Lin et al., 2013); *orco*^2^ has a null mutation in the *orco* gene (Larsson et al., 2004); *gr63a*^1^ has a null mutation in the *gr63a* gene (Jones et al., 2007); *or42b*^EY14886^ has an insertional mutation that disrupts the *or42b* gene (Bellen et al., 2004); *shakB*^2^ was from R. J. Wyman and carries a nonsense mutation in the signal sequence of shakB, rendering it a functional null (Krishnan et al., 1993); *ir25a*^1^ has a null mutation in the *ir25a* gene (Benton et al., 2009); *UAS-opGCaMP6f* and *lexAop-opGCaMP6f* were a gift from Barret D. Pfeiffer and David J. Anderson and were used for all functional calcium imaging experiments; *UAS-CsChrimson-mVenus* expresses a Venus-tagged red-shifted channelrhodopsin (Klapoetke et al., 2014) under Gal4 control. *UAS-DTI* and *lexAop-DTI*, which express the DTI mutant form of diphtheria toxin subunit A in a Gal4 (Han et al., 2000)- or lexA-dependent manner, respectively, were used for cell ablations.

*lexAop-DTI* flies were generated in this study by first PCR amplifying the *DTI* gene from the *pUAS-DTI* plasmid (Han et al., 2000), a gift from Leslie M. Stevens, and replacing (at the ATG) the open reading frame for *myr∷GFP* in plasmid *pJFRC19-13xlexAop2-IVS-myr∷GFP* (Pfeiffer et al., 2010) with that of *DTI* using isothermal assembly (Gibson et al., 2009). *pJFRC19-13xlexAop2-IVS-myr∷GFP* was from Gerald Rubin (Addgene plasmid #26224). Site-specific transgenic flies were generated using standard methods (Bestgene, Inc., Chino Hills, CA) to insert the *lexAop-DTI* transgene into the attP2 landing site.

#### Odor Delivery

Odors were delivered essentially as previously described (Hong and Wilson, 2015). The olfactometers used for odor stimulus delivery are diagrammed in Figure S1. Briefly, for all odors except CO_2_, a constant stream of air (200 ml/min) was directed at the fly. Ten percent of the airstream (20 ml/min) was routed through the “normally open” port of a three-way solenoid valve (ASCO 411-L-1324-HVS, NISCO, Inc., Duluth, GA) and passed through the headspace of a control vial filled with solvent, before rejoining the main carrier stream. When triggered by an external valve command, the three-way valve redirected the 20 ml/min odor stream to exit through the “normally closed” port and into the headspace of the vial filled with odor, thereby further diluting the concentration in the headspace by 10-fold prior to delivery to the fly. The 20 ml/min control or odor streams were carried by tubing of matched lengths and rejoined the carrier stream at the same point along the carrier tube, approximately 10 cm from the fly. Odor concentrations are reported as v/v dilution in paraffin oil (J.T. Baker, VWR #JTS894), with the exceptions of ammonia and acetic acid, which were diluted in water. The headspace for all odors was diluted by a factor of 10 in air. The only exceptions to this were acetic acid, which was always diluted 2-fold, and ammonia, which was diluted 100-fold in Figure 1, in air.

For delivery of CO_2_, the olfactometer was modified such that the odor stimulus line and the control balance line were each controlled by their own 3-way solenoid valve. The stimulus line exited from the “normally closed” port of its valve (the “normally open” port of that valve was vented outside the microscope box). The balance line exited from the “normally open” port of its valve (the “normally closed” port of that valve was vented outside the microscope box). Flow rates in the stimulus and balance lines were always equal, and stimulus line and balance line valves each received the same external command signal. The stimulus and balance lines each joined a constant carrier stream of air (180 ml/min) at the same point along the carrier tube (Figure S1), approximately 10-cm from the end of the tube. In this way, the total flow rate experienced by the fly was kept constant. For instance, for the delivery of a 5% CO_2_ stimulus, the olfactometer functioned as follows. Under default (non-stimulus) conditions, the balance line, carrying 10 ml/min of air, joined the carrier stream, carrying 190 ml/min of air, to direct a total of 200 ml/min of air at the fly; the 10 ml/min of 100% CO_2_ in the stimulus line was vented outside. When the command signal was high, the stimulus line, carrying 10 ml/min of 100% CO_2_, joined the carrier stream, carrying 190 ml/min of air, to direct a total of 200 ml/min of 5% CO_2_ at the fly; the 10 ml/min of air in the balance line was vented outside. In this way, CO_2_ stimuli of varying concentration were generated by adjusting the relative flow rates of the carrier line (from 180 to 199 ml/min), and the stimulus and balance lines (from 1-20 ml/min).

For either olfactometer, the opening of the carrier tube measured 4 mm in diameter and was positioned approximately 1 cm from the fly. A small funnel (10-cm diameter) connected to a vacuum line was placed behind the fly to vent odors. Flow rates were metered using mass flow controllers (MC series, Alicat Scientific, Tucson, AZ), and all air was first passed through a charcoal filter before being routed into the mass flow controllers. A total flow rate of 200 ml/min at the fly was used for all experiments, except for the optogenetic experiments in Figure 4E-G and the experiments in Figure 6, where a total flow rate of 2 L/min was used. In Figure 4E-G, the flow rate was increased to reduce the latency of the odor stimulus to better match the very short latency of the light stimulus. In Figure 6, a higher flow rate was used to reduce low-pass filtering of rapidly fluctuating odor stimuli.

#### Two-photon calcium imaging

In vivo functional calcium imaging was performed essentially as previously described (Hong and Wilson, 2015). The fly was head-fixed, the dorsal cuticle was removed, and the antennal lobes were exposed. The antennae were snugly secured below the imaging chamber, keeping them dry and allowing them to remain responsive to odors. Antennal lobes were imaged from the dorsal side; horizontal imaging planes were acquired at varying depths along the dorsal-ventral axis of the antennal lobes.

GCaMP6f fluorescence was excited with 925 nm light from a Mai Tai DeepSee laser (Spectra-Physics, Santa Clara, CA). Images were acquired with an Olympus 20X/1.0 numerical aperture objective (XLUMPLFLN20XW) on a two-photon microscope equipped with galvo-galvo scanners (Thorlabs Imaging Systems, Sterling, VA) at 5.5 frames s^-1^ at a resolution of 224×224 pixels covering an area of 90 × 90 μm^2^ (Figures 1 and 2) or 140 × 140 μm^2^ (all other data). The collection filter was centered at 525 nm with a 50 nm bandwidth, with the exception of the optogenetic experiments (see below). The microscope was housed in a lightproof box, and experiments were conducted at room temperature (∼22°C). The brain was constantly perfused by gravity flow with saline containing (in mM): 103 NaCl, 3 KCl, 5 N-Tris(hydroxymethyl)methyl-2-aminoethane-sulfonic acid, 8 trehalose, 10 glucose, 26 NaHCO_3_, 1 NaH_2_PO_4_, 1.5 CaCl_2_, and 4 MgCl_2_ (pH 7.3, osmolarity adjusted to 270– 275 mOsm). The saline was bubbled with 95% O_2_/5% CO_2_ and circulated in the bath at ∼2-3 ml min^-1^.

Specific glomeruli were identified using a combination of their anatomical depth, location, size, and shape in the baseline fluorescence signal, which are invariant across flies. Their identity was confirmed by evaluating their characteristic odor response properties, using a test panel of odors presented at relatively low concentration (spanning 10^-4^ to 10^-8^). These odors sparsely activate ORNs that express the highest affinity odorant receptors for the odor (Hallem and Carlson, 2006; Hong and Wilson, 2015), reliably identifying the glomerul(i) to which they project. For experiments in Figure 1 where we surveyed for glomeruli that are responsive to CO_2_, the antennal lobe of many flies (>10 flies) was systematically sampled at 5μm intervals all along the entire depth of the antennal lobe while repeatedly presenting CO_2_ to the fly. Glomeruli exhibiting reliable CO_2_-evoked calcium activity across individuals were then identified using the above criteria.

Imaging trials were each of 30 s duration, with the stimulus delivered 10s after the onset of imaging. The response to a given stimulus was measured as a block of three replicate trials, and stimulus blocks were delivered in pseudo-random order. The intertrial interval was 3 s. For experiments in Figures 3C-D; 4, 5, and Figure S5, recordings from experimental and control flies were interleaved. When collecting odor-evoked responses for each stimulus-glomerulus pair, the average amount of time elapsed from the start of the experiment to the measurement was roughly matched between genotypes. This matching was done to ensure that any differences observed between genotypes was not due to systematic differences in the time course of the modulation of CO_2_-evoked responses in VA2 ORNs. The only exception to this was for a small subset of *gr63a*^-/-^ flies, from which we recorded CO_2_-evoked responses in VA2 ORNs two hours after the start of the experiment, to account for the possibility that the mutation may simply delay the conversion of the response. Excitatory responses to CO_2_ were never observed, although responses of VA2 ORNs to diacetyl were present.

#### Optogenetics

For the experiments in Figure 4E-G, flies were raised on cornmeal/molasses food supplemented with one teaspoon of potato flakes rehydrated 1:1 (v/v) with 140 μM all-trans retinal in water. All-trans-retinal was prepared as a 35 mM stock in ethanol and stored at −20°C. The parental cross that generated experimental flies was carried out in the dark on all-trans-retinal-supplemented medium, and newly eclosed experimental flies were maintained in the dark, also on all-trans-retinal medium, until used in experiments.

Calcium imaging of optogenetically evoked signals was performed essentially as described above with the following modifications. The collection filter for imaging was centered at 500 nm with a 20 nm bandwidth (HPX500-20, Newport Corporation, Irvine, CA). The use of this filter allowed simultaneous detection of GCaMP6f fluorescence emission while stimulating with red light, without stimulus bleed-through to the detector. Light was delivered from the tip of an optical fiber (400-μm core, 0.39 NA, Thorlabs, Newton, NJ) butt-coupled to a 625nm LED (M625F2, 1000 mA, Thorlabs, Newton, NJ). The tip of the optical fiber was positioned ∼ 1 mm away from the antennae, which were snugly tucked beneath the stainless steel floor of the imaging chamber.

Odor and light stimulus trials were interleaved during the experiment. A 1 s pulse of light was used for optogenetic stimulation. This pulse duration was chosen based on the results of pilot single-sensillum recordings of light-evoked firing rates in ab1C ORNs in the flies (which express CsChrimson in ab1C ORNs and GCaMP6f in *orco*-positive ORNs, Figure 4E). Light pulses of greater than 1 s duration resulted in ORNs firing a short burst of spikes (∼250 ms) and subsequently entering into depolarization block.

#### Pharmacology

All drugs were bath applied in the saline. Mecamylamine was used at 100 μM. Picrotoxin and CGP54626 were used at 5 μM and 50 μM, respectively. Oubain was used at 100 μM. Drugs were washed-in for 5 min before initiating recordings.

#### Single-sensillum Recordings

Single-sensillum recordings were performed essentially as previously described (Bhandawat et al., 2007). Briefly, flies were immobilized in the end of a trimmed pipette tip, and one antenna was visualized under a 50x air objective. The antenna was stabilized by tightly sandwiching it between a set of two fine glass hooks, fashioned by gently heating pipettes pulled from glass capillaries (World Precision Instruments, TW150F-3). A reference electrode filled with saline (see above) was inserted into the eye, and a sharp saline-filled glass recording microelectrode was inserted into the base of the sensillum. ab1sensilla were identified based on their characteristic size and morphology, position on the antenna, and the presence of four distinct spike waveforms (in wild type flies), each having a characteristic odor sensitivity (de Bruyne et al., 2001) (see also Figure S3). Signals were acquired with a MultiClamp 700B amplifier, low-pass filtered at 2 kHz, and digitized at 10 kHz. Delivery of odor- and light-stimuli was carefully matched to that in functional imaging experiments. To unambiguously distinguish ab1A from ab1B spikes in some critical experiments, it was necessary to kill one of these ORN types using diphtheria toxin expression (Figure S3A-B and Figures 3A-B and 5A-B).

#### Measuring CO_2_-evoked neural responses

CO_2_-evoked responses in ORNs in glomeruliVA2 and DM1 generally converted over the course of a recording from a “low” state, dominated by inhibition, to a “high” state, characterized by mixed excitation and inhibition (Figure 2). In separate experiments, we found that the modulation of the CO_2_ response does not require prior odor exposure and can be observed with just two presentations of CO_2_, one measured immediately after the fly is placed on the recording rig and one measured thirty minutes later (data not shown).

The fact that the time course of the modulation of the CO_2_ response could vary significantly across different flies (Figure S2) presented a challenge to defining the “early” and “late” response. In a typical experiment, including the experiments in Figure 2A-C, the “early” response of all glomeruli of interest to all stimuli of interest was measured within the first five minutes after placing the fly on the rig. The response of ORNs in VA2 was then periodically probed by delivering CO_2_ to the fly every ∼10 min, until an excitatory response was observed, plateaued in amplitude over consecutive measurements, and at least 30 min had elapsed since the start of the experiment. At this point, the response of all glomeruli to all stimuli was measured again and defined as the late response. Rarely, a fly responded to CO_2_ with a mixed excitatory/inhibitory response on the very first presentation of CO_2_ (Figure S2B, green trace). More typically, the duration of the full conversion of the CO_2_ response ranged from ∼10-60 min. The CO_2_ response in VA2 ORNs converted to the “high” state in almost all flies. A similar protocol was used for measuring the modulation of the response to acetic acid in VA2 ORNs (Figure 2D). In Figures 1 and 3-6, whenever possible, responses were collected from preparations that achieved the conditions we have defined as the late response; if the experimental genotype did not allow this, responses were collected at a time >30 minutes from the start of the recording. When comparing between paired control and experimental genotypes (e.g., Figures 3 and 4), the average amount of time elapsed from the start of the experiment to the measurement of each glomerulus-stimulus pair was roughly matched between genotypes (see above).

CO_2_-evoked responses in ORN terminals in glomerulus DM1 also change; although we have not studied it systematically, we have observed that the modulation of CO_2_ responses in DM1 ORNs does not necessarily occur concurrently with that in VA2 ORNs.

We also considered the hypothesis that the modulation of CO_2_-evoked calcium signals in VA2 ORN terminals originates from spiking activity in ab1B ORNs, which project to VA2. Thus, we investigated whether the response of ab1B ORNs to CO_2_ might change over the duration of an experiment. In the sensillum recordings in Figure 3A, flies were mounted and positioned on the rig for >60 min prior to recording. We implemented this waiting period to allow for any potential conversion of ab1B responses to CO_2_; however, ab1B ORNs were never observed to spike in response to CO_2_. These experiments suggest that the modulation of CO_2_-evoked calcium signals in VA2 ORN terminals does not arise from changes in ab1B spiking activity, which is also consistent with other experiments demonstrating that they have a lateral origin.

#### Data analysis

##### Two photon functional imaging

Imaging analysis was performed in MATLAB (Mathworks, Natick, MA) using custom scripts. Regions of interest (ROIs) defining specific glomeruli were manually drawn in each imaging plane from the movie of raw fluorescence signal, using anatomical position, size, shape, and odor tuning criteria, as described above. Calcium transients (ΔF/F) were measured as changes in fluorescence (ΔF) normalized to the mean fluorescence during the baseline period (F, averaged in the 10 s prior to stimulus onset). All imaging data was background subtracted (using the mean pixel intensity outside the antennal lobe) prior to analysis. In each experiment, the calcium response in an ROI was computed as the mean across three trials of each stimulus. Unless otherwise indicated, the mean calcium responses reported in the figures are averaged across independent replicates of each experiment and represent the mean ± s.e.m. computed across flies. The *n* for each figure are in Table S1.

To generate heatmaps of peak responses in individual flies, ΔF/F was calculated on a pixel-by-pixel basis, and three consecutive frames centered on the peak of the response were averaged. For each stimulus, data were pooled by averaging the peak odor-evoked heat map (*Δ*F/F) across three trials. A Gaussian low-pass filter of size 5 × 5 pixels was applied to *Δ*F/F maps.

##### Single-sensillum recordings

Spikes were detected using custom scripts in MATLAB. Peristimulus time histograms (PSTHs) were computed by counting the number of spikes in 50-ms bins that overlapped by 25 ms. Single-trial PSTHs were baseline subtracted (using the spontaneous firing rate in the pre-stimulus period) and averaged together across five trials for each stimulus to generate the PSTH describing the response to an odor in a given experiment. The mean PSTHs shown in the Figures 3B and 5B, D, F represent the mean ± s.e.m. computed across multiple flies. The *n* for each figure panel are listed in Table S1.

### Statistical Analysis

The number of replicates for each condition in each experiment is reported in Table S1. The number *n* represents the number of individual flies in which each measurement was made. The specific statistical test used and its results are reported in the figure legends. Statistics were computed in Matlab or Prism 8 (GraphPad Software, San Diego, CA). Reported *p*-values were corrected for multiple comparisons within a given experiments. Results that were not significant (n.s.) had *p-*values greater than *α*=0.05.

---

**Table S1:**
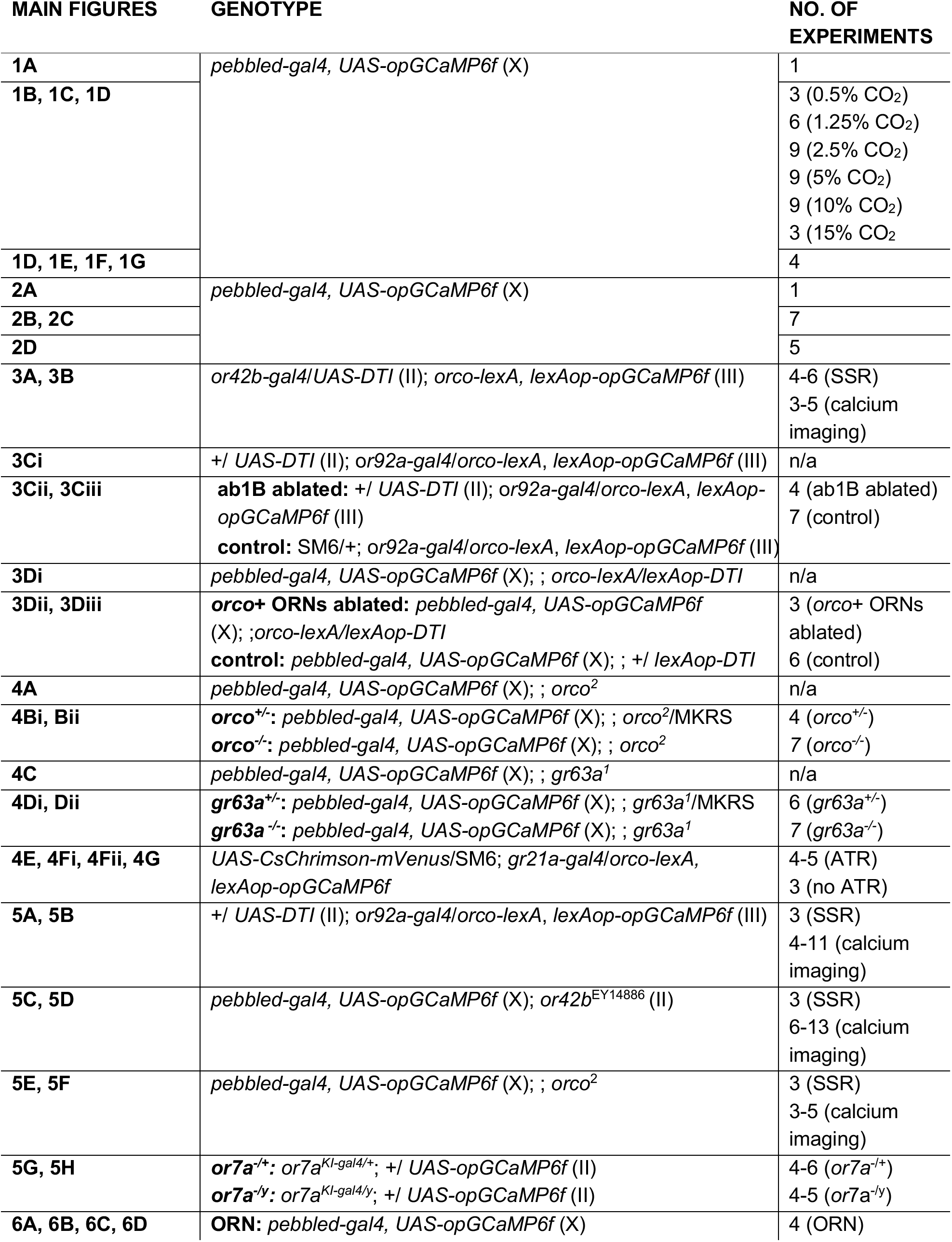
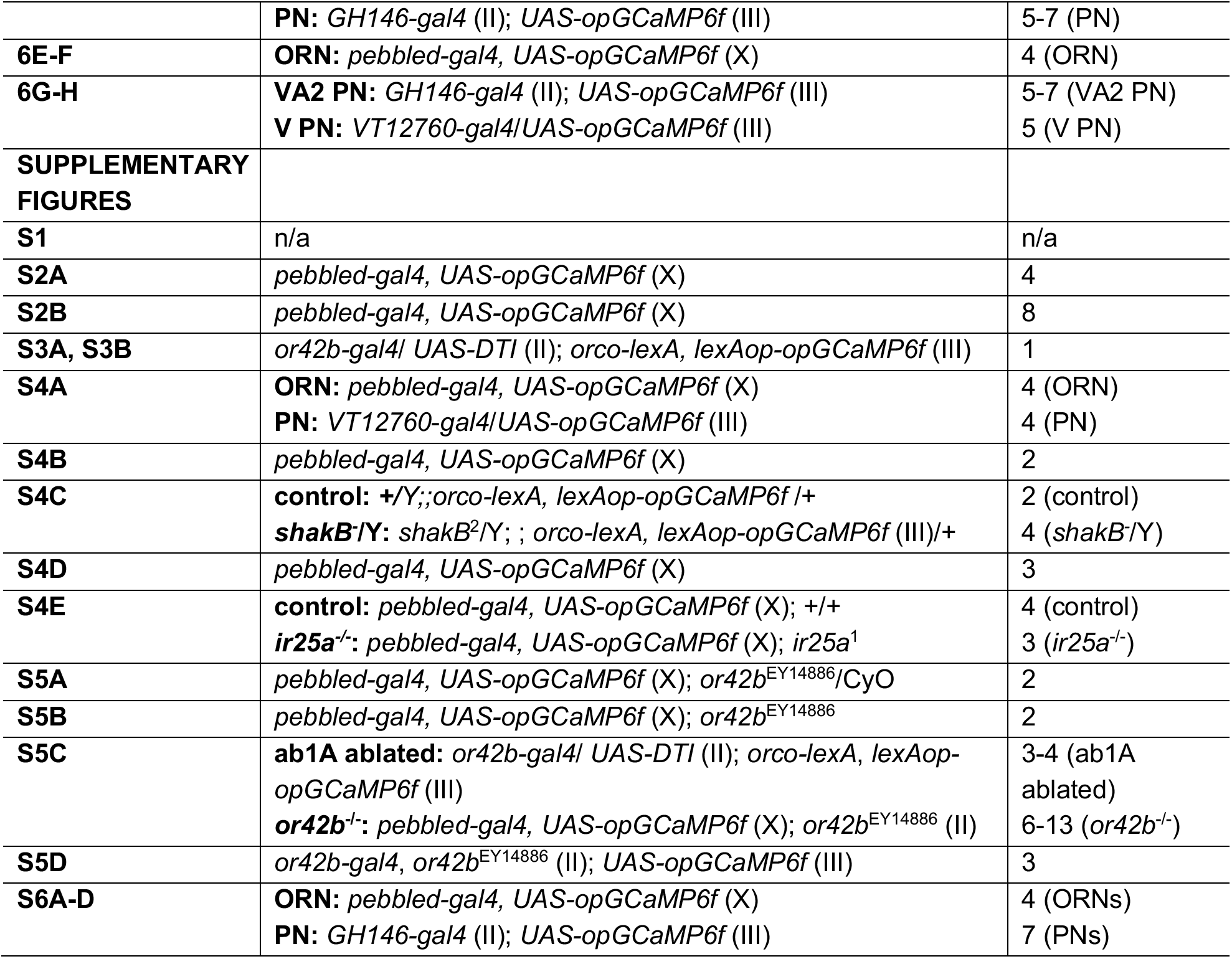
Complete genotypes and n for all experiments.

